# Vpr shapes the proviral landscape and polyclonal HIV-1 reactivation patterns in cultured cells

**DOI:** 10.1101/2021.10.08.463623

**Authors:** Edmond Atindaana, Sarah Emery, Cleo Burnett, Jake Pitcher, Jeffrey M. Kidd, Alice Telesnitsky

## Abstract

Cell culture models suggest that the HIV-1 viral protein R (Vpr) is dispensable for latency establishment. However, whether Vpr affects the persistent proviral landscape and responsiveness to latency reversing agents (LRAs) is unclear. Here, integration site landscape, clonal dynamics, and latency reversal effects of Vpr were studied by comparing barcoded *vpr+* and *vpr-* populations arising after infection of Jurkat cells in vitro. The results showed that individual integrant clones differed in fractions of LTR-active daughter cells: some clones gave rise to few to no LTR-active cells while for others almost all daughter cells were LTR-active. Integrant clones with at least 60% LTR-active cells (high LTR-active clones) contained proviruses positioned closer to preexisting enhancers (H3K27ac) and promoters (H3K4me3) than clones with <30% LTR-active cells (low LTR-active clones). Comparing *vpr*+ and *vpr*- populations revealed that the *vpr+* population was depleted of high LTR-active clones. Complementing *vpr*-defective proviruses by transduction with *vpr* 16 days after infection led to rapid loss of high LTR-active clones, indicating that the effect of Vpr on proviral populations occurs post-integration. Comparing *vpr*+ and *vpr*- integration sites revealed that predominant *vpr*+ proviruses were farther from enhancers and promoters. Correspondingly, distances to these marks among previously reported intact HIV proviruses in ART-suppressed patients were more similar to those in the *vpr+* pool than to *vpr-* integrants. To compare latency reactivation agent (LRA) responsiveness, the LRAs prostratin and JQ1 were applied separately or in combination. *vpr*+ and *vpr*- population-wide trends were similar, but combination treatment reduced virion release in a subset of *vpr*- clones relative to when JQ1 was applied separately, an effect not observed in *vpr+* pools. Together, these observations highlight the importance of Vpr to proviral population dynamics, integration site landscapes, and responsiveness to latency reversing agents.

**One Sentence Summary**

Expression properties and responsiveness to latency reactivation agents of individual HIV-1 proviral clones within polyclonal populations are masked by dominant clones and influenced by proviral proximity to certain epigenetic marks and by Vpr, a viral factor not previously known to affect latency and reactivation.

## Introduction

HIV-1 establishes stable, long-lived reservoirs in patients treated with highly active antiretroviral therapy (HAART). Within this reservoir, cells that contain transcriptionally silent replication competent proviruses are inert to immune recognition and clearance as well as to current antiretroviral therapies; thus, even optimally treated patients remain infected (1). A cure for HIV- 1 will require either ridding an individual of all infected cells (sterilizing cure) or reducing the reservoir to levels that enable the patient’s immune control (functional cure).

Previous studies have demonstrated that latency establishment and maintenance is multifaceted and can involve the depletion of cellular factors such as the transcription factor NFkB, reduction in the level of the positive transcription elongation factor b (pTEFb), sequestration of pTEFb by the HEXIM/7SK RNA complex, the lack of CDK9 phosphorylation by CDK7 to promote pTEFb assembly, and promoter occlusion due to transcriptional interference from upstream host genes (2–7). The remodeling of the Nuc-1 nucleosome within the long terminal repeat (LTR) promoter, modulation of chromatin accessibility by histone modifying enzymes, and/or methylation of CpG sequences can also play roles in establishing and maintaining HIV-1 latency (8–10). Viral proteins and genetic elements may also contribute to latency, including deficiency of the viral transactivating protein Tat, which recruits pTEFb to the HIV promoter, deficiency of Rev, which mediates the export of unspliced HIV-1 RNA to the cytosol, or mutations in TAR or RRE *cis* acting RNA elements, which regulate these functions (11, 12).

Cure strategies that are under active investigation include “shock and kill” and “block and lock”. In contrast to HAART, which prevents new infection, the shock and kill method involves inducing virus expression, with the intention that this will lead to either cytopathic death of reactivated cells or host immune recognition and clearance (13). Some proposed approaches involve perturbing cellular pathways with small molecules or stimuli that complement intracellular deficiencies and can reverse HIV-1 latency in experimental models. For example, prostratin and phorbol 12- myristate 13-acetate (PMA) can reactivate latent HIV-1 through activation of protein kinase C (PKC), leading to an increase in the level of NFkB. Other latency-reversing agents (LRAs) include those which act to increase the level of pTEFb, including the BET bromodomains inhibitor JQ1 and a class of drugs known as ingenols. JQ1 notably inhibits the bromodomain of BRD4 and prevents BRD4 from binding acelylated histones at promoters and specific enhancers, leading to a decrease in lineage specific genes (14). Additional classes of LRAs act to modify the chromatin environment: suberoyl anilide hydroxamic acid (SAHA) and entinostat being members of the class that promotes a euchromatin state by inhibiting histone deacetylases (15–19).

The size of a patient’s latent reservoir is estimated to be between one in a million and three in ten thousand resting memory CD4+ T cells (20). The rarity of latently infected cells in virally suppressed individuals makes the study of *in vitro* latency models necessary. Tissue culture models developed to study HIV-1 latency and reactivation differ from one another in that some utilize infectious virus while others employ HIV-1-based reporters. Prominent among this later class are systems like the widely used J-Lat clones, which assess LTR promoter activity by reporter gene expression but lack HIV-1 genetic elements believed to be dispensable in HIV-1 latency (21–23). One such element is the *vpr* gene, whose product has been shown to play roles in efficient viral infectivity, enhances CCR5-tropic HIV-1 replication in humanized mice, acts as a coactivator of the LTR and enhances HIV-1 transcription by Vpr-mediated ubiquitination and degradation of Phosphorylated CTD Interacting Factor I (PCIFI) (24–27). Vpr also causes cell-cycle arrest and induces cell death through the depletion of the chromosome periphery protein CCDC137, and induces widespread changes in host gene expression in CD4+ T cells (28–30). For these reasons, some latency models use *vpr*-defective proviruses to allow the survival of cells harboring transcriptionally active proviruses. Cultured over time to allow proviral transcriptional silencing, these cells can then be used for reactivation studies (31–34).

Results from LRA reactivation studies using these models have been inconsistent, as demonstrated by Spina et al.(35). This discordance may be due to differences in the models of latency used (22). Some approaches used to produce silenced proviruses exclude latent clones that are established shortly after infection (23, 36). Moreover, HIV-1’s semi-random integration site preference makes it likely that in each model, the integrants in a particular clonal cell line being studied have unique host chromosomal loci that may be subject to locus-specific regulation. The extent to which integration sites influence the reactivation of latent proviruses has long been recognized but not extensively explored (32, 37).

Molecules that are effective at reversing latency in various tissue culture models have been identified, but evidence thus far suggests that these are either too toxic to be therapeutically useful or that they fail to reduce the size of the latent reservoir in patients (19, 38–41). Fundamental aspects of latency in patients, such as the size of the latent reservoir, remain poorly defined. Recent studies have demonstrated that quantitative viral outgrowth assays (QVOA) underestimate the latent reservoir while PCR-based quantitation can overestimate it due to the predominance of defective proviruses (20), leading to ongoing efforts to improve reservoir quantification (42, 43).

Since accurately measuring the latent reservoir is challenging in and of itself, it seems likely that measuring changes in the latent reservoir before and after LRA treatment will be equally challenging. Because HIV-1 latency can be established and maintained by multiple mechanisms (4), an effective therapy will likely be one that disrupts multiple pathways.

To date, there is limited evidence addressing whether or not mechanistically distinct classes of LRAs affect all proviruses uniformly or have differential effects on individual proviral clones. In this study, the clonal structure and influence of hundreds to thousands of individual proviruses’ host genomic loci on reactivation dynamics were investigated, with responses to distinct classes of LRAs in latent *vpr-* and *vpr+* proviral pools compared. The results revealed differences in the clonal structures of populations that influenced total levels of reactivation and showed that the aggregate properties of polyclonal populations misrepresented the behavior of individual proviruses. Whether the presence of *vpr* affected integration at the time of infection or if its subsequent expression altered the structure of polyclonal populations or influenced activities of residual cells was examined. In response to dual treatment with JQ1, which affects pTEFb availability, and prostratin, which activates PKC, individual cell clones within polyclonal *vpr-* provirus populations exhibited either enhanced, additive or repressed reactivation, while the spectra of responses were more limited in type and magnitude in cell clones containing *vpr+* proviruses. Together these findings suggest that Vpr plays a critical role in shaping transcriptionally inactive proviral populations and in defining their LRA responsiveness.

## Results

### vpr- and vpr+ proviral pools differed in numbers of LTR-active cells

To study the effects of Vpr on the responsiveness of individual proviruses in the absence of virus spread, Jurkat cells were infected with distinguishable *vpr+* or *vpr-* versions of the NL4-3 derivative shown in Fig. 1A (44). Using the EF1α promoter to drive constitutive expression of the puromycin resistance gene allowed selection of infected cells independent of LTR expression and thus the retention of proviruses silenced shortly after infection. Each genomic RNA in the infecting virus contained a unique 20b randomized “barcode” inserted in U3, which was duplicated in both LTRs after reverse transcription and enabled tracking individual proviruses. We refer to the barcodes as “zip codes” because in the context of proviruses, they report the genetic neighborhoods of each integrant. Infected cell populations were passaged without cell cloning, thus generating polyclonal integrant populations, within which transcriptionally active cells were identified using LTR-driven GFP expression or by progeny genomic RNA in released Env- virions.

**Figure 1.**
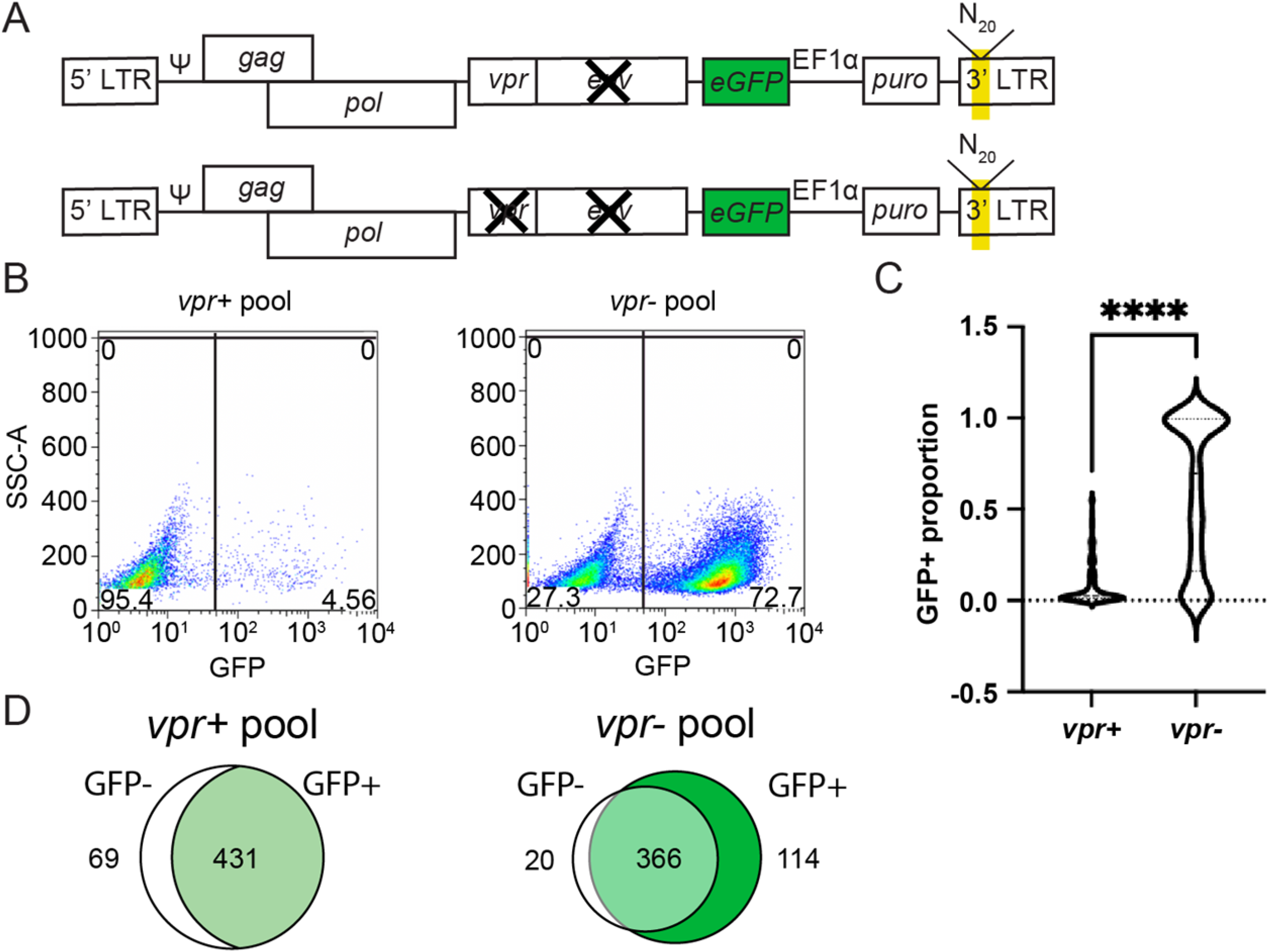
Generation of zip coded HIV-1 proviral pools. Polyclonal cell pools containing *vpr+* and *vpr-* proviruses were established in Jurkat T cells and compared. **A.** Schematic of vectors used to produce zip coded virus (not to scale). Vectors were pseudotyped by co-transfection with pHEF-VSV-G into HEK293T cells. The yellow shade in the 3’ LTR indicates position of 20bp randomized barcode in U3. **B.** Representative flow cytometry plots of polyclonal Jurkat T cells generated by infection with barcoded *vpr+* (left panel) or *vpr-* (right panel) viruses, that were puromycin-selected for four days and expanded for an additional ten days. X axis indicates GFP signal detected in the FITC channel; numbers in each gate indicate percent of cells gated GFP- or GFP+. **C.** A comparison of the numbers of high and low LTR-active clones in *vpr+* and *vpr-* pools. The fractions of total cells in each clone that sorted GFP+ (LTR-active) were calculated for the most abundant 500 clones in each pool by quantifying the fraction of each zip code (integrated barcode) in the GFP+ sorted cells’ genomic DNA to its quantity in the unsorted pool by high throughput sequencing and normalizing for the percent of the unsorted pool that was GFP+. Violin plots compare the GFP+ proportions (indicative of clones’ fraction of LTR-active daughter cells) for the top 500 most abundant zip codes each pool. (p= **** indicated shows p<0.0001 Mann-Whitney U two-tailed test.). **D.** Venn diagrams of the number of zip codes in GFP+ and GFP- sorted sub pools for *vpr+* and *vpr-* pools. The number in the white region indicates the number of zip codes observed only in GFP- sorted cells, that in the light-green region of intersection represents the number of zip codes that were present in both GFP+ and GFP- sorted cells, and the number in the dark green region represents zip codes present in only the green sorted cells. Reported zip codes represent those present in the top 78% and 88% of reads for *vpr+* and *vpr-* respectively when clones were ordered by read abundance.

Previous work using zip coded *vpr-* derivatives of this vector has shown that for each integrant, the clone gives rise to a mixture of cellular progeny that includes some GFP+ cells and some GFP- cells (44), and that sibling cells switch between LTR-active (GFP+) and -inactive (GFP-) expression phenotypes in a phenomenon we called “flickering”. To confirm that individual integrant clones in the current study also contained clone-specific mixtures of GFP+ and GFP- cells, single clones were isolated by limiting dilution from the *vpr*- pool and expanded, and cells from each clone were then subjected to flow cytometry. Consistent with previously reported differences in bimodal expression patterns among proviral clones, the results showed that each clonal pool was comprised of both GFP+ and GFP- cells, with GFP+ proportions that differed among the clones: mostly GFP+ cells for some clones, and distinctly different GFP+ proportions for others (S1B Fig.).

In the current study, four independent polyclonal integrant pools were established by infecting roughly 5x10^6^ Jurkat T cells at a multiplicity of infection of <0.0005 to ensure infected cells contained one provirus per cell. Two of the pools contained *vpr+* proviruses and two had integrants lacking *vpr*. Flow cytometry analysis after fourteen days of expansion showed significantly fewer GFP+ cells in the two *vpr*+ infected cell pools than in the two *vpr-* provirus pools (p=0.042, paired t-test; <5% vs. 73% GFP+, respectively Fig. 1B, S1A Fig.). These fourteen days post-infection cell pools were sorted into GFP+ and GFP- subpools, cellular DNA was harvested from an aliquot of the sorted cells immediately after sorting, and the zip codes of integrants were amplified from the cellular DNA and catalogued by high throughput sequencing.

After analyzing roughly three million sequencing reads per pool, zip codes were ordered by read abundance. Determining how many unique barcodes were present in each GFP+ pool revealed that similar *numbers* of zip codes (approximately 2000)—each indicative of a single integration event—were detected in the GFP+ sorted cells from both *vpr+* and *vpr-* pools, even though the GFP+ fraction of total cells in the *vpr+* pool was very low. This finding suggesting that the number of GFP+ integrants initially established did not differ markedly between pools is consistent with previous work in dendritic cells that indicated that the extent of proviral integration does not differ depending on the presence or absence of Vpr (45).

Combining population-wide GFP+ fractions in the unsorted cells with zip code read counts within sorted sub-pool libraries allowed calculating the proportion of LTR-active cells within each cell clone (Fig. 1C). Notably, for the top 500 most abundant clones in the unfractionated *vpr+* and *vpr-* pools, median GFP+ proportions were significantly lower in the *vpr+* pool (p=0.0001, Mann-Whitney U two-tailed test), indicating that most cells in the *vpr+* pool were members of low LTR-active clones (Fig. 1C). When GFP+ sorted cells from *vpr*+ and *vpr*- pools were cultured further, no viable *vpr*+ GFP+ cells were detected after three days, and thus their clonal structures could not be analyzed further. These results indicated that integration events that resulted in GFP expression were equally likely in *vpr*+ and *vpr*- proviral pools when the cells were examined early post-infection. However, the depletion of the Vpr+ GFP+ cell sub-populations suggested that whenever cells flickered “on”, the resulting GFP+ cells died upon further cell culturing.

### Vpr shapes the clonal structure of infected pools

Observing fewer GFP+ cells in the *vpr*+ pool than in cells with *vpr-* proviruses was not surprising due to Vpr’s well-known cytotoxic effects. However, a small fraction of *vpr+* GFP+ cells were observed among unsorted *vpr+* cells, even though sorted *vpr+* GFP+ cells did not survive three days of culture.

One plausible reason unsorted *vpr+* populations contained rare GFP+ cells was that these cells arose via recent phenotypic switches from clones that were largely inactive. To test this possibility, zip codes in GFP+ and GFP- sorted subpools were compared for both *vpr+* and *vpr-* integrants (Fig. 1D). Ordering zip codes by their abundance and analyzing those that comprised the top 500 revealed that among *vpr-* cells, about 73.2% of all the zip codes were found in both subpools, while 4% were observed only in GFP- cells and 22.8% only in the GFP+ subpool. In contrast, 86.2% of the *vpr+* cells’ zip codes were found in both subpools, with 13.8% observed only in the GFP- subpool, and none of the zip codes found exclusively in the GFP+ subpool. This suggested that the small fraction of GFP+ cells in the *vpr+* pool (Fig. 1B; left panel) resulted from the recent acquisition of LTR expression by cells from the larger GFP- cell pool. If the *vpr-* pool is assumed to be relatively representative of a population that results when all clones are equally viable, this suggests that when initially integrated, proviruses whose daughter cells were largely or always GFP+ were the dominant class of clones.

### High LTR-active clones’ proviruses are in close proximity to genome marks associated with active gene expression

The finding that LTR-active *vpr+* cells were rapidly lost suggested that proviruses that had integrated into more active regions of the host genome might be selected against within polyclonal *vpr+* populations. Therefore, we compared the proximities of *vpr+* and *vpr-* integration sites to genomic features associated with active gene expression. First, zip code integration sites were determined using cellular DNA harvested ten days after establishment of each pool and mapped to 1171 and 1121unique sites in the human genome for *vpr+* and *vpr-* respectively (see Methods). Then, the proportion of zip codes located in genes versus those not in genes (as defined by ENCODE for Jurkat cells (46) were compared (Fig. 2A, S2 Fig.). The results indicated that similar majorities of integrants were established within genes regardless of whether or not *vpr* was present. Next, *vpr+* and *vpr-* integrants were compared for their proximity to specific genome marks associated with active gene expression that have been reported to pre-exist in Jurkat cells (47). No differences were found in distances to closest DNase I sensitivity sites, which are associated with open chromatin (Mann-Whitney U test, p=0.1854) (Fig. 2B). However, proximities to H3K27ac (associated with enhancers) (Mann-Whitney U two-tailed test, p<0.0001) and H3K4me3 (associated with active promoters) (Mann-Whitney U two-tailed test, p<0.0001) marks differed significantly, with proviruses from *vpr+* pools farther from these marks (Fig. 2C-D).

**Figure 2.**
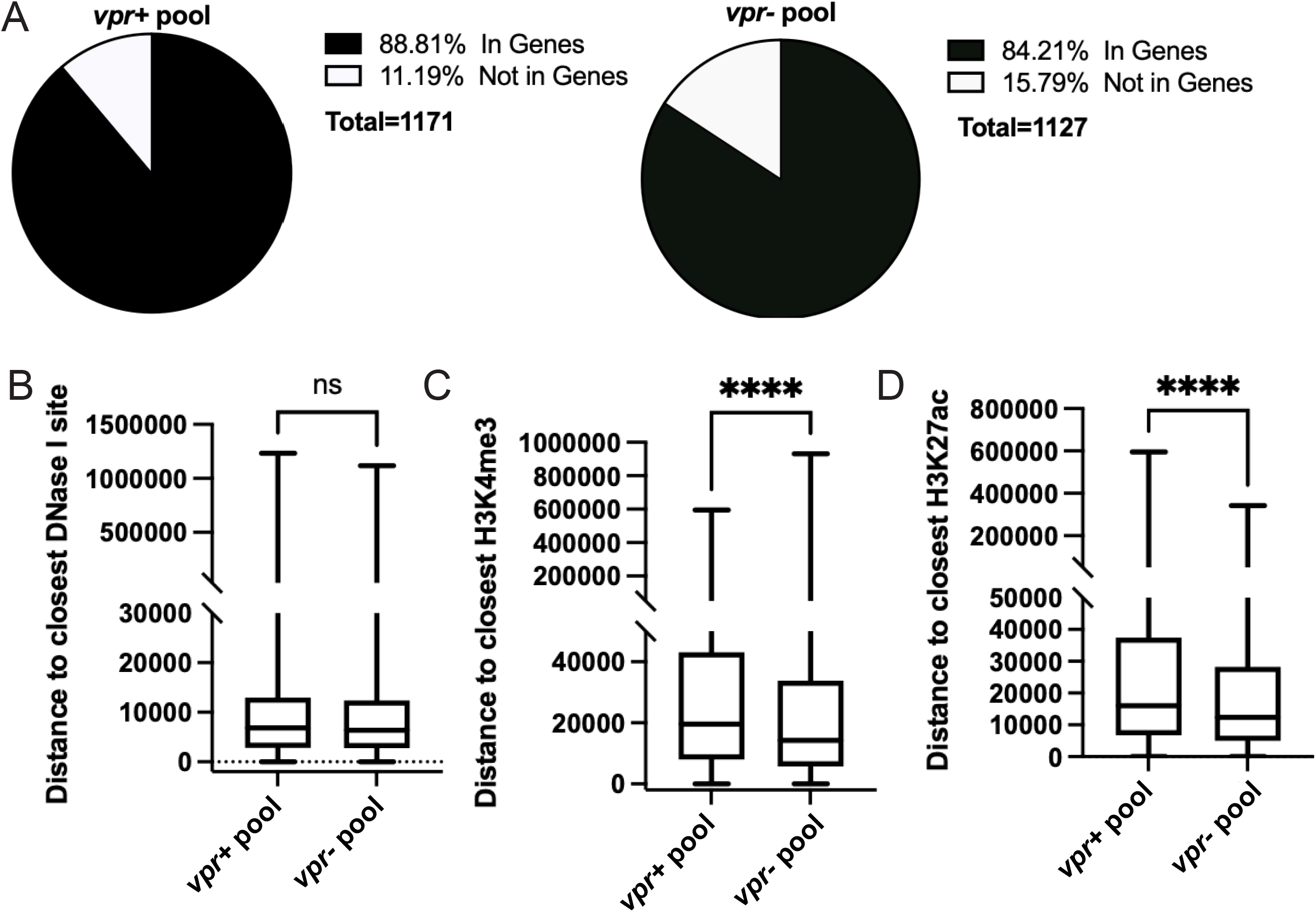
Comparison of integration sites in *vpr+* and *vpr-* pools. Integration sites for both *vpr+* and *vpr-* pool cells harvested 10 days post-infection were determined and distances to the nearest genome features indicated were mapped and compared. A. Pie charts showing the percentage of integration sites found within genes and those not found in genes for *vpr+* and *vpr-* pools. B.-D. Box plots showing pairwise comparisons between *vpr+* and *vpr-* integration sites’ distances to the closest: B. DNase I hypersensitivity site, C. H3K4me3, and D. H3K27ac. Mann-Whitney U two- tailed test was conducted for pairwise comparisons for B-D (p= ns, and **** indicated shows p>0.05 and p<0.0001 respectively).

Next, integrants from *vpr-* pool were grouped into high LTR-active clones (GFP+ proportion >= 60%) and low LTR-active clones (GFP+ proportion < 30%) and compared for their proximities to H3K27ac and H3K4me3 (Fig. 3A). The results showed that high LTR-active clones were closer to both marks than low LTR-active clones and suggested that the difference in integration site proximity observed between *vpr+* and *vpr-* proviruses is due to the survival of high LTR-active clones in the absence of Vpr.

**Figure 3.**
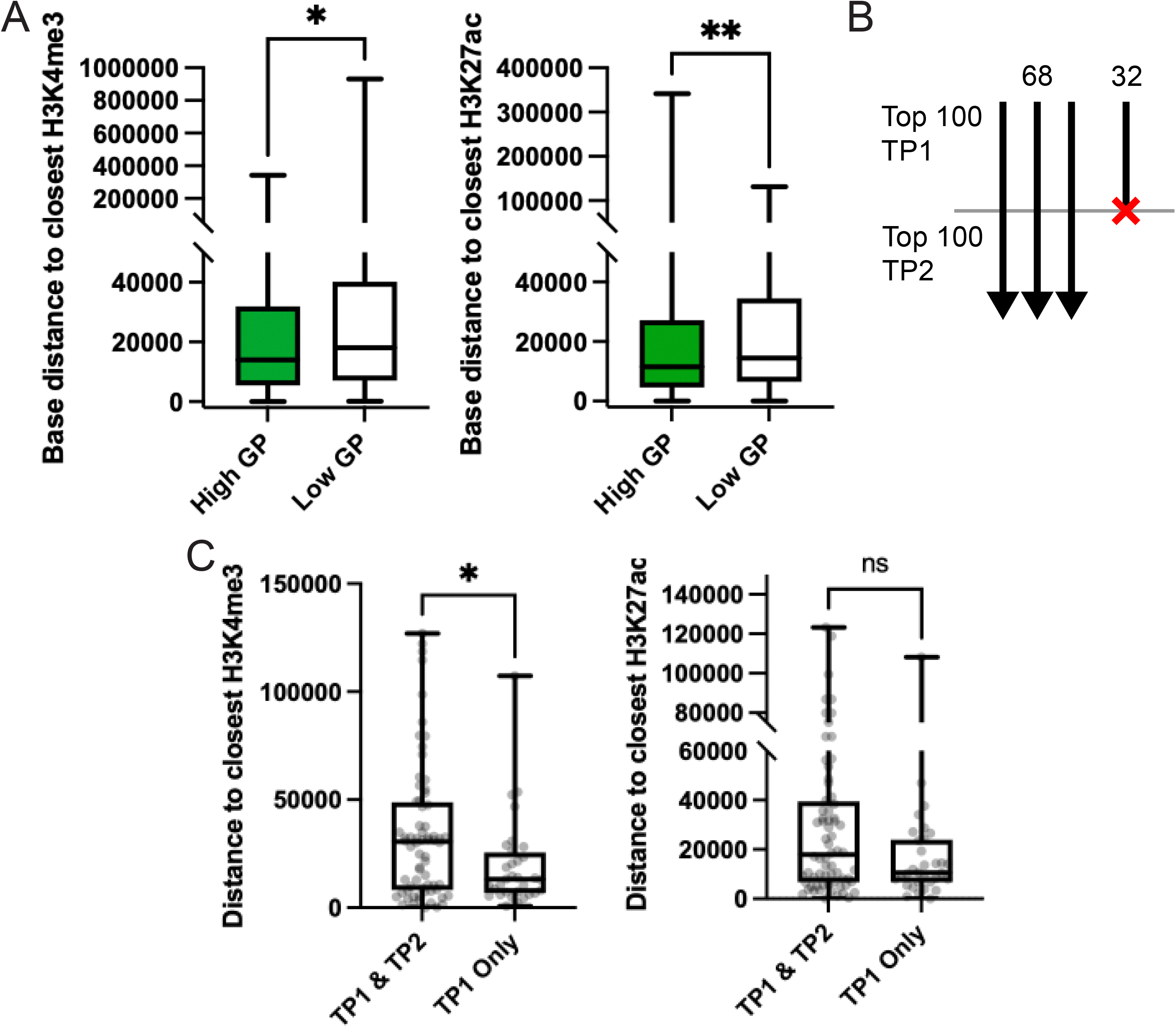
Integration site comparison within *vpr+* and *vpr-* pools. A. The GFP+ proportion of each zip code from the *vpr-* pool was determined by quantification of the abundance in GFP+ cells’ genomic DNA to the total abundance in the genomic DNA from unsorted cells. Zip codes with GFP+ proportion >= 0.6 were binned as “High” and zip codes (595 zip codes) with less than 0.3 GFP+ proportions were binned as “Low” (222 zip codes). Pairwise comparison of the distance to the closest H3K4me3 (left) and H3K27ac (right) marks. B. Schematic diagram showing changes in *vpr+* pools’ clonal composition over time. The number of zip codes in the top 100 most abundant clones of the unsorted *vpr+* pool sampled at time point one (TP1; 10 days post-infection) and the number of zip codes that remained in the top 100 clones at time point two (TP2: 24 days post- infection) are indicated. Arrows from TP1 to TP2 indicates zip codes present at both time points. Arrows with red crosses indicate zip codes that were no longer present in the top 100 clones. C. Box plots comparing integration site distances to active chromatin marks for the *vpr+* pools over time. The closest H3K4me3 or H3K27ac mark for zip codes that were among the top 100 clones at both TP1 &TP2 and for the clones that were lost from the top 100 at TP2 are indicated (p=ns, indicated shows p>0.05; Mann-Whitney U two-tailed test).

Because the *vpr+* pool was devoid of high LTR-active clones, its high- and low LTR-active integrants could not readily be compared. However, with the reasoning that abundances of residual LTR-active clones would gradually diminish over time, the dynamics of the v*pr+* pool were examined by comparing samples harvested two weeks apart (Fig. 3B). Examination of the 100 most abundant zip codes in the *vpr+* cell pool showed that only 68 of the top 100 zip codes on day 10 post-infection (time point 1, or TP1) were observed among the 100 most abundant zip codes on day 24. When integration site proximities to H3K4me3 and H3K27ac marks of these zip codes were compared, those that were absent from the top-100 on day-24 (time point 2, TP2) (Fig. 3C) were significantly closer to H3K4me3 (p=0.0193, Mann-Whitney U two-tailed test), and tended to be closer to H3K27ac marks (p=0.1581, Mann-Whitney U two-tailed test) than the TP 1 clones that remained within the top-100 most abundant clones at TP2.

### The spectra of LRA responses among vpr+ and vpr- clones differed

Next, the reactivation properties of *vpr-* and *vpr+* pools were compared and the behaviors of individual proviral clones within the populations were determined. The latency reactivation agents (LRAs) prostratin, a protein kinase C agonist, and JQ1, a bromodomain inhibitor, were applied separately or in combination.

To examine reactivation in the total LTR-inactive cell sub-population of each pool, FACS-sorted GFP- cells from both *vpr+* and *vpr-* pools were subjected to prostratin (10 μM), JQ1 (2 μM), or both drugs in combination for 24 hours. Reactivation was measured both by changes in the frequency of GFP+ cells using flow cytometry and by the amount of virus release (Fig. 4A-B; S3A-B Fig.). The results indicated that compared to single prostratin and JQ1 treatments, dual treatment resulted in additive levels of reactivation in both *vpr+* and *vpr-* populations by both criteria compared to single prostratin or JQ1 treatment (Fig 4A shows reactivation monitored by GFP+ cells and Fig. 4B shows virus release; left panel indicates reactivation for *vpr+* and right panel for *vpr-*). In dual LRA treatments for both pools, there was an approximately 4-fold increase in GFP+ cells, while virus release increased by approximately 30-fold relative to DMSO control. The most significant difference between *vpr-* and *vpr+* pools was that the absolute amount of virus release was 3-fold higher in *vpr-* pools, and the responsiveness to JQ1 was lower in the *vpr+* pools. The differences between pools in their extents of reactivation were not due to differences in cell viability (S3C Fig.). The observation that reactivation was enhanced by dual prostratin and JQ1 treatment is consistent with previous works by Boehm *et al.,* and Darcis *et al.,* using the same drugs in cell culture models of latency and ex vivo treated cells from HIV patients respectively (21, 48).

**Figure 4.**
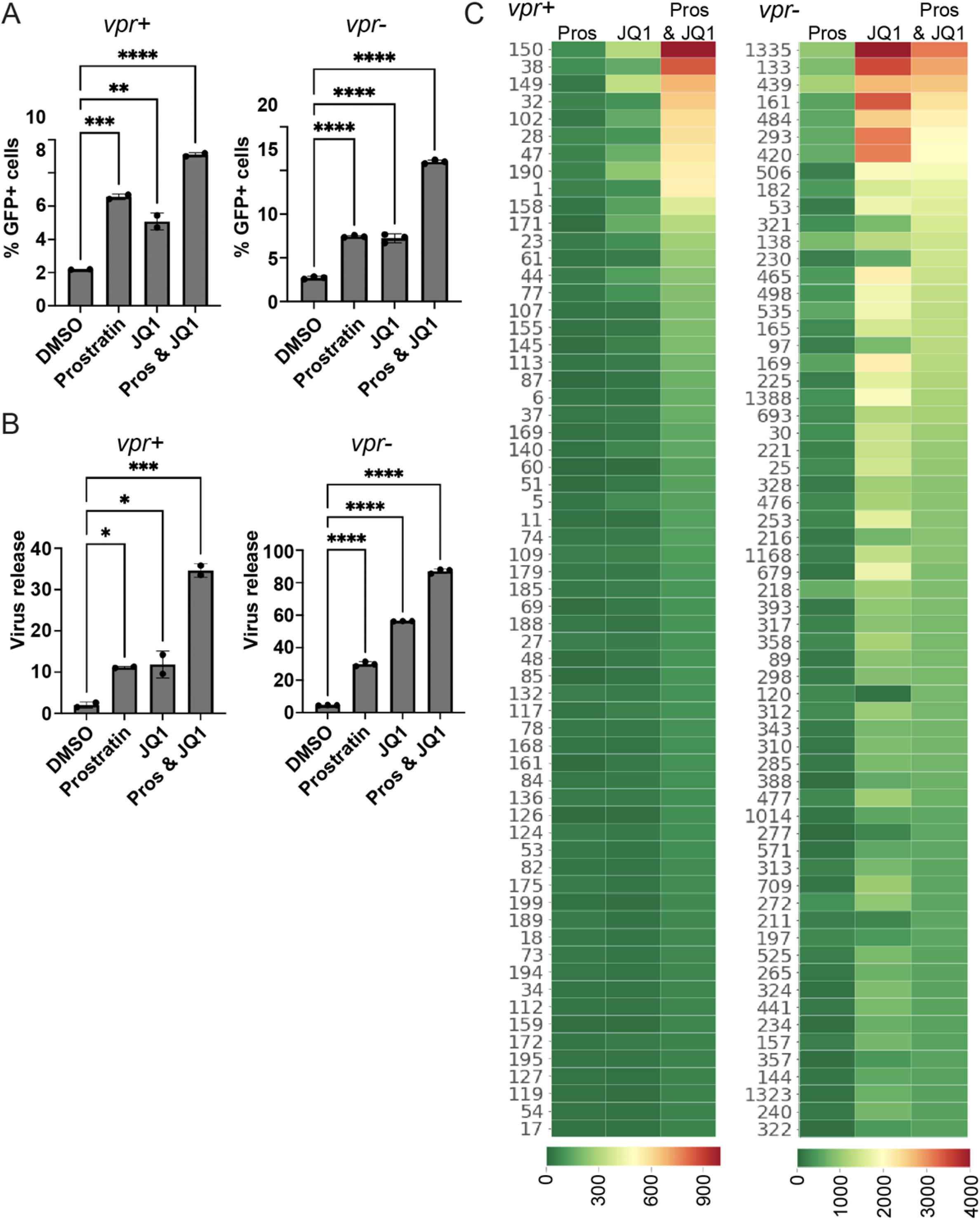
Effects of LRAs on zip coded pools and clones. *vpr+* and *vpr-* GFP- cell fractions were exposed to 0.1% DMSO, 2µM JQ1, 2µM prostratin, or to a combination of prostratin and JQ1. Reactivation to GFP+ was measured at 24 hours post-infection by flow cytometry, and virus release was quantified using a reverse transcriptase assay and normalized to define p24 levels. A. Bar graphs showing the frequency of GFP+ cells post LRA treatment (left panel for each pool) and B. the amount of virus released into culture supernatant (right panel for each pool) for the indicated polyclonal pools. The error bars show the mean and standard deviation from two experimental treatment repetitions. C. Heatmaps of the clonal (zip code) virus release per treated GFP- cell (left: *vpr+* pool and right: *vpr-* pool). Numbers at the left of each panel are clone identifiers generated by ordering proviral zip codes in decreasing relative abundance, as determined for the unsorted pools. Every row represents a unique cell clone’s response. The clones were ordered from top to bottom by diminishing virus release per treated cell upon dual LRA treatment. The color bar indicates the extent of release per treated cell in arbitrary units.

To investigate the effects of the treatments on individual GFP- clones, total virus release was quantified by p24 equivalent and cDNA was generated using viral genomic RNA upon LRA treatment. Zip codes were amplified from the viral cDNA and also from an untreated aliquot of the GFP- cellular DNA and high through-put sequenced. The results were normalized to calculate average virus release per treated cell for each clone (Fig. 4C; S3D Fig.). Analysis revealed that when cells were treated with both drugs, some clones in both the *vpr+* and *vpr-* pools displayed enhanced virus release per treated cell compared to single treatment conditions. Interestingly, both proviral pools included a subset of clones that were not detectably reactivated by either prostratin or JQ1 when the LRAs were applied alone, but that were reactivated upon dual LRA treatment. Surprisingly, and only in the *vpr-* pool, reactivation levels observed under dual LRA conditions were lower than those observed with single LRA use for a subset of clones (Fig 4C: right panel. Compare, for example, *vpr-* JQ1 and Pros +JQ1 columns). These same patterns of reactivation were observed when experiments were repeated using a second set of independently established *vpr-* and *vpr+* pools (S3D Fig.).

The finding that some clones were detectably reactivated by dual LRA treatment but not by either prostratin or JQ1 alone was interesting. A possible explanation for this was that these clones were in low abundance. If rare clones released little virus when prostratin and JQ1 were applied separately, their release might only rise to a detectable level in the cDNA library when these drugs were combined. However, the results showed that these were not rare clones. For example, the most abundant clone in the GFP- *vpr-* library (pool 2; clone number 3), which accounted for about 15% of the treated GFP- pool cells (Supplementary Table), was one of the clones for which detectable virus was released only upon dual LRA treatment.

### Complementation of the vpr-defective pool led to depletion of high LTR-active clones

As an additional test of the effects of Vpr on polyclonal population composition, a functional copy of the *vpr* gene was added to cells harboring *vpr-* proviruses 16 days post-infection. This was achieved by transducing the *vpr-* pool with a *tat*-deficient lentiviral vector containing LTR driven *vpr,* which also contained the fluorescent marker *mKO* expressed from the constitutive SV40 promoter. This resulted in expression of Vpr only when a pre-existing *vpr-* provirus was transcriptionally active. mKO expression was used as a proxy for *vpr* vector transduction, and enabled cells that were successfully transduced to be sorted in the phycoerythrin (PE) channel (Fig. 5A). 48 hours post-transduction, an unsorted aliquot was saved while the remaining cells were sorted to identify GFP- PE+ cells (S4 Fig.A-B) for work described below.

**Figure 5.**
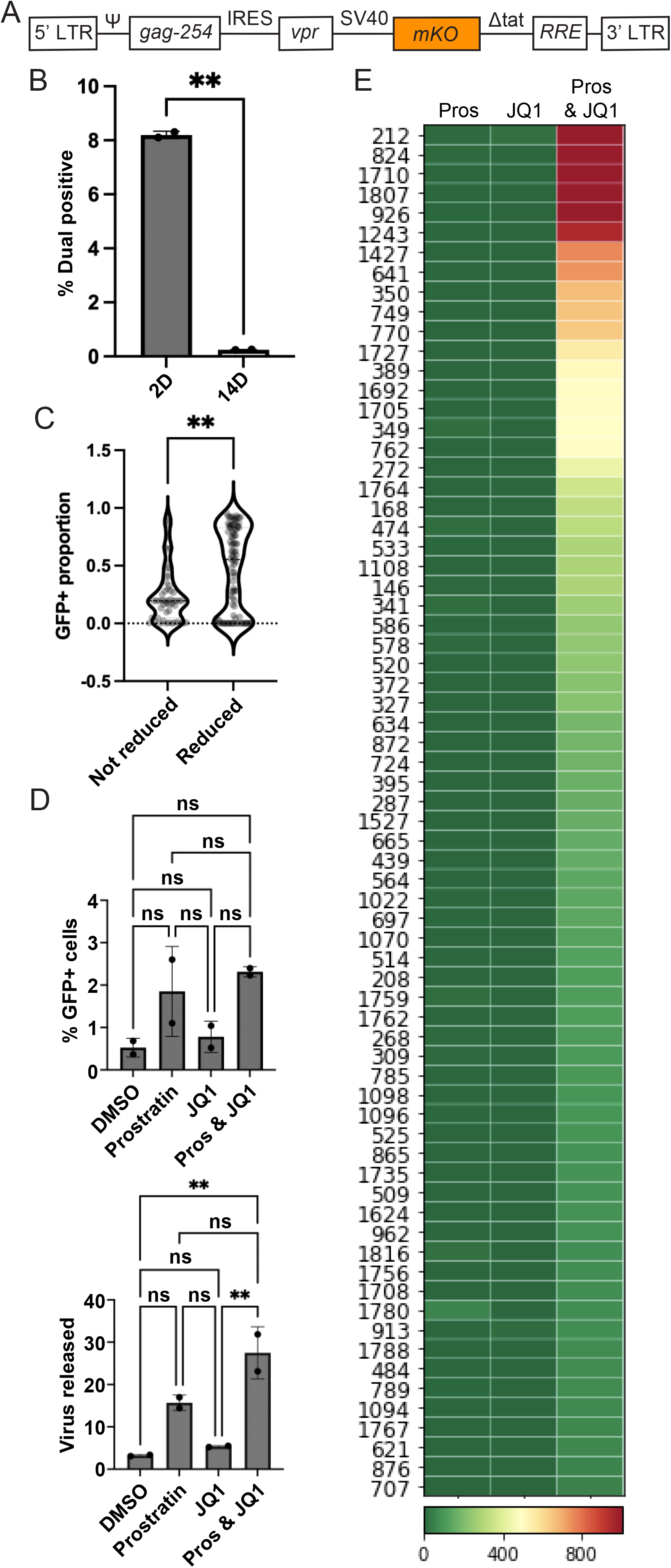
Complementation of *vpr-* pools with Vpr expression vector. *vpr* was added to *vpr-* pools and effects on clone sizes were determined by high throughput analysis. A. Schematic diagram of the Tat-deficient Vpr expression vector (not to scale), which contains *IRES-vpr* under the HIV-1 LTR promoter and an SV40 promoter-driven mKO fluorescent reporter. B. Bar graph of the percent GFP+ cells for two *vpr*^rev^ pools established in parallel at day-2 (2d) and day-14 (14d) post-transduction with the Tat-deficient Vpr expression vector (p=0.0076, significant pairwise comparison by paired t-test). C. Pairwise comparisons of GFP+ proportions between the clones that were reduced (10-fold or more reduction in relative abundance) and those that were not reduced (less than 1.5-fold reduction in relative abundance) after *vpr* addition (significant pairwise comparison by Mann-Whitney U two-tailed test). D. Bar graphs showing the percent of GFP+ cells 24 hours post reactivation with JQ1, prostratin, and dual prostratin-JQ1 for the *vpr^rev^* pool (left) and *vpr-* pool (right). E. Heatmap of the clonal (zip code) virus release per treated GFP- cell of *vpr^rev^* pool. Numbers at the left of each panel are clone identifiers generated by ordering proviral zip codes in decreasing relative abundance, as determined for the unsorted pools. Every row represents a unique cell clone’s response. The clones were ordered from top to bottom by diminishing virus release per treated cell upon dual LRA treatment. The color bar indicates the extent of release per treated cell in arbitrary units.

Comparing the proportions of GFP+ cells in the parental *vpr-* pool to the proportions in the same pool “reverted” *to vpr+* by introduction of the *vpr* vector (hereafter referred to as the *vpr^rev^* pool) showed a marked decrease in GFP+ cells between day-2 and day-14 post transduction (Fig. 5B). This selective loss of GFP+ transduced cells corroborated observations with the *vpr+* pool that LTR-active cells were depleted in the presence of Vpr.

Proviral zip code abundances in unsorted *vpr-* and *vpr^rev^* pools were then compared. Of the 500 most abundant clones in the *vpr-* pool, zip codes were split into two groups based on observed fold-change in relative abundance: those that were reduced by 10-fold or more in *vpr^rev^* (reduced clones), and those with no observed reduction in their relative abundance upon *vpr* addition (not reduced). A comparison between the “reduced” and “not reduced” groups revealed that nearly all *vpr-* pool clones that displayed high GFP+ proportions--that is, clones in which most member cells displayed LTR activity-- were reduced in the *vpr^rev^* pool (Fig. 5C).

The *vpr^rev^* pool was then used to address whether *vpr-* pools rendered *vpr+* by complementation displayed reactivation patterns similar to those of the original *vpr+* pools. Transduced GFP- *vpr^rev^* subpools were subjected to prostratin, JQ1, or dual prostratin-JQ1 treatment and analyzed by flow cytometry and virus release (Fig. 5D). In general, the magnitude of reactivation in the transduced cells was diminished across all treatment conditions relative to parental *vpr-* GFP- cells, with cells responding less to JQ1 treatment than they did before Vpr addition. Consistent with expectations if the phenotypes above reflected Vpr, transduced cells showed enhanced reactivation upon dual LRA treatment (Fig. 5E; S4C Fig.). Notably, clones (such as clones numbered 53, 133, 161, 293, 420, etc.) that were highly responsive to JQ1 treatment in the parental untransduced *vpr-* pool, and that differentiated the LRA responsiveness of the *vpr-* pool from the *vpr+* pool, were highly represented among those lost upon Vpr addition.

### Genomic features of persistent intact proviruses in patients on ART are more similar to vpr+ than vpr- pool members

The proviruses in HIV patients on ART contain intact Vpr and have survived its selective pressures. Having observed differences between *vpr+* and *vpr-* viruses in cultured cells in terms of their reactivation patterns and proximity to genome features, we sought to determine which class of these proviruses more closely resembled those in persistent clinical isolates.

To do this, previously published data on patients’ intact provirus integration sites (49) were analyzed for their proximity to H3K4me3, H3K27ac, and DNase I hypersensitivity sites that have been reported for Jurkat T cells and primary CD4+ T cells (see Methods) (47). When these proximities were compared to those in the *vpr-* and *vpr+* pools established here, prominent proviruses in the *vpr-* pool, and not those in the *vpr+* pool, were found to significantly differ from the patient proviruses in their proximities to H3K4me3 and H3K27ac marks (Fig 6. A-B; S5 Fig.), but not DNase I hypersensitivity sites (Fig. 6C). These results suggest that the integration site distribution of persistent proviruses in patients across the tested genome marks was similar to those observed in *in vitro* established *vpr+* proviral populations but significantly different from what was observed within *vpr-* proviral pools.

**Figure 6.**
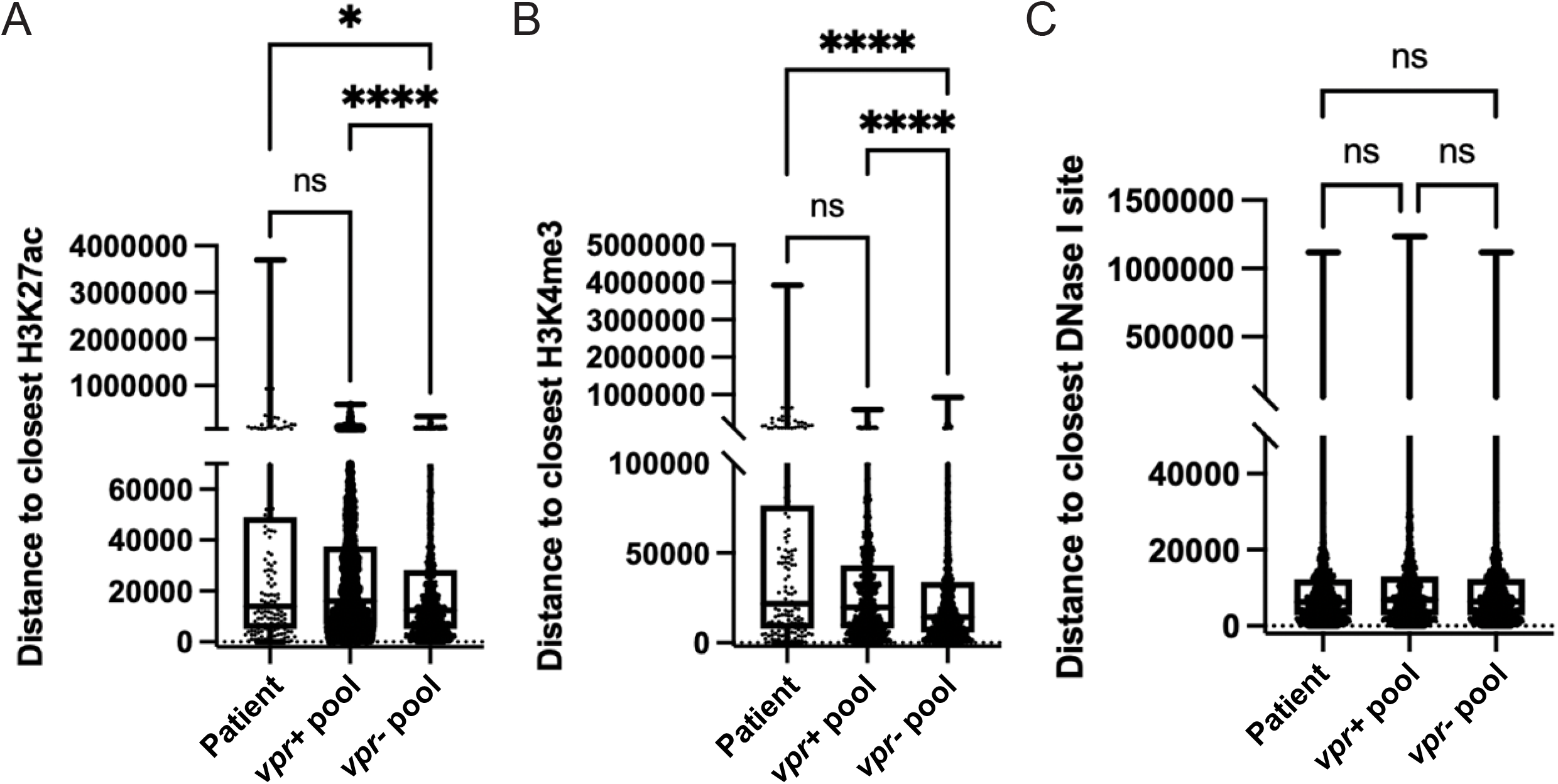
Comparison of patients’ integration site features with *vpr+* and *vpr-* pools. Previously published HIV integration sites from three patients were used to determine base distances to closest H3K27ac, H3K4me3, and DNase I sensitivity sites. Distances were compared to those of *vpr+* and *vpr-* pools. Box plots comparing base distances of *vpr+*, *vpr-*, and patients’ proviral integration sites to: A. H3K27ac, B. H3K4me3, and C. DNase I sensitivity sites. (Kruskal- Wallis test. p=ns, *, **, ***, **** indicate p>0.05, < 0.05, <0.01, 0.001, and 0.0001 respectively).

## Discussion

This work describes how Vpr shapes the overall provirus population structure, dominant host integration site landscape, and responsiveness to latency reversing agents in cultured cells. Cell-based models are critical to latency reactivation agent drug discovery and other HIV persistence work, but these models differ by cell type, form of HIV-1 used, whether they use isolated clones or polyclonal populations, and other parameters. Studies using polyclonal pools may implicitly assume that the pools contain an equivalent proportion of each proviral clone. However, as demonstrated here and previously, even when clonal target cell lines are infected, pools rapidly become dominated by a subset of proviral clones (44). Thus, population-wide trends may not apply to all clones. Similarly, because proviral clones differ in expression patterns, clonal latency models’ HIV-1 expression properties may represent the biases of the investigators who chose the clones more than selective pressures that shape the latent reservoir.

Primary CD4+ T cells are the closest tissue culture system to natural infection since CD4+ T cells are the primary target of HIV infection. However, these cells only survive 2-4 weeks *ex vivo* unless treated with antiapoptotic agents (38, 50) and do not proliferate unless stimulated. We previously compared expression patterns of zip coded proviruses in primary CD4+ T cells to those in Jurkat T cells, and observed indistinguishable spectra of expression patterns (44). However, extended cell passaging was required in the current work. Thus, we chose to use Jurkat cells, which limits the physiologic relevance of our findings, although Vpr has been noted to exert similar effects in primary CD4+ and Jurkat T cells (51).

Because identifying latent cells is challenging, many experimental latency systems include reporter genes and/or delete viral genes believed to be unnecessary for silencing and reactivation (33, 34, 52). For example, two prominent studies that used barcoded proviruses to track virus dissemination in animals used *vpr-* proviruses (33, 52). The impact of this variation is not well understood.

The zip coded provirus approach used here enabled addressing Vpr effects by comparing *vpr+* and *vpr-* pools. *vpr+* pools were found to contain significantly fewer GFP+ cells (indicative of transcriptionally active proviruses) than did *vpr-* pools, but similar numbers of proviral clones were detectable in both pools soon after their establishment. Consistent with previous findings (44), for some *vpr-* integrant cell clones, essentially all member cells displayed HIV gene expression (which we designated high LTR-active clones); for other clones most cells did not express HIV (low LTR-active clones), and a smaller number of clones displayed similar numbers of active and inactive member cells. In contrast, within the most abundant *vpr+* clones, most clonal member cells did not express HIV genes (low LTR-active clones). These findings may help explain why estimates of the transcriptionally active fraction of HIV-1 infected cells vary among previous studies. Using polyclonal *vpr+* proviruses *in vitro*, Dahabieh *et al*. suggested that most integrated proviruses are latent, even though it has been estimated that latent cells represent only a small fraction of total infected cells in patients (29, 36, 53). Consistent with Dahabieh *et al.,* we found that HIV is transcriptionally inactive in most infected cells that persisted in *vpr+* populations, but our additional work suggested this reflected the selective survival of low LTR-active clones.

Previous work has shown that within proviral clones, individual cells can shift from being silenced to expressing their proviruses, while maintaining the overall proportions of their daughter cells that possess active proviruses (37, 44, 54). For example, when barcodes are used to examine *vpr-* proviral clones within polyclonal infected cell populations, individual member cells within each clone either express HIV genes, as monitored by GFP reporter, p24 staining, or virion release, or they do not express HIV. Although some diminution of LTR expression is observed over time, for the most part each clone adopts a stable, heritable pattern of bimodal gene expression (44). The current work showed *vpr*+ and *vpr*- polyclonal populations did not differ upon integration but differed significantly in proportion of LTR-active cells after passaging. Together, these observations suggest that the low proportion of transcriptionally active cells in *vpr+* integrant pools might reflect the selective depletion of high LTR-active clones (Fig.7); a notion that is further supported by observations in *vpr^rev^* pools, which were *vpr-* pools that were complemented with a Vpr expression vector. According to this model, although the proportion of total cells that were GFP+ was significantly lower in polyclonal *vpr+* integrant populations than in *vpr-* proviral pools at 14 days, the spectrum of *vpr+* integration sites was likely indistinguishable from *vpr-* clones upon initial establishment.

**Figure 7.**
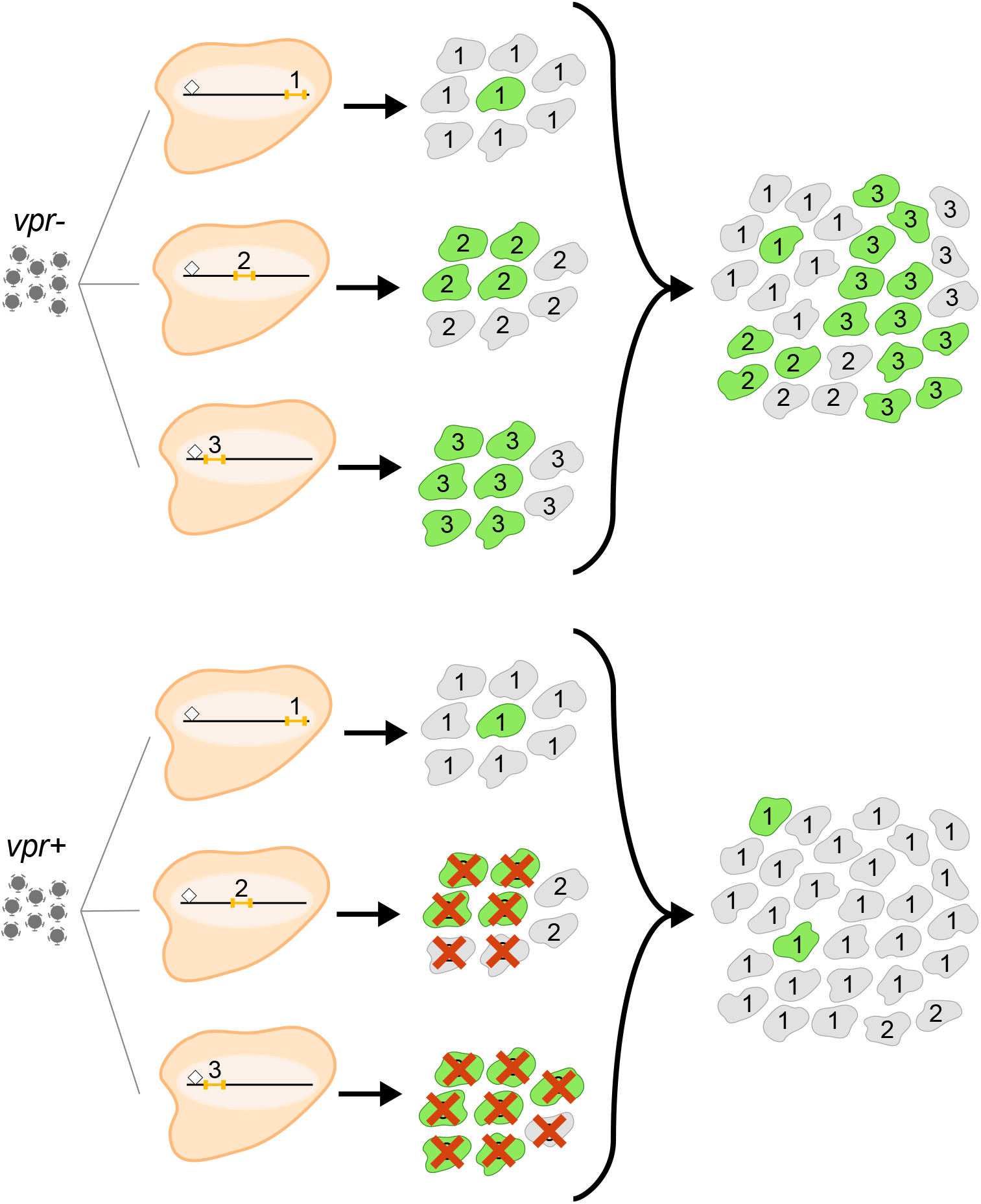
Schematic model of the effect of Vpr on integrant populations’ proviral landscapes. From left: proviruses are integrated at the same genomic positions (designated 1, 2, and 3 above the small orange bars, which indicate proviruses) regardless of whether the infecting virus was Vpr+ or Vpr-. Note that integrants 1, 2, and 3 differ in their distances to the closest chromatin mark, which is indicated by a small diamond, which may affect their expression characteristics and responses to LRAs. Next, infected cells divide to form cell clones. For clone 1, cells that express HIV genes (indicated by green cell) are rare (low LTR-active clone) and for other clones, more cellular members of each clone express HIV (eg: clone 3: high LTR-active clone). This pattern of bimodal gene expression is an intrinsic property of each clone. The red Xs indicate that high LTR-active clone members are selectively depleted by Vpr. As a result, in the polyclonal populations shown at right, low LTR-active clones are enriched in the *vpr+* pool.

A little-discussed problem with transcriptionally silenced provirus models is that they tend to “spontaneously revert” during passage and generate small proportions of cells with transcriptionally active proviruses (37, 55). Our previous work and findings here suggest an explanation slightly different from “spontaneous reversion” for the appearance of LTR-active cells in passaged latency model cells. As evident from quantifying phenotypic switching within expression-sorted polyclonal subpopulations, the phenotypes of active- and inactive- subpopulations of each clone are not static, but instead equilibrate in a phenomenon we call flickering. This suggests most LTR-inactive cells are not stably silenced but rather represent the transient off-state of an on-off continuum. If so, “spontaneous reversion” by a small subset of latency model cells, like the low proportion of GFP+ cells in the current study’s *vpr+* pool, might more accurately be viewed as the programmed equilibration of a low LTR-active clone than the outlier behavior in a subset of silenced cells.

Comparing integrants that dominated passaged *vpr+* and *vpr-* pools suggested Vpr selects against proviruses proximal to genome marks associated with active gene expression. Consistent with previous work, *vpr+* and *vpr-* pools showed no difference in preference for integration into actively transcribed genes (56–58). However, predominant *vpr+* proviruses, which represented clones with the largest numbers of daughter cells, were farther from the closest enhancer or promotor than were predominant *vpr-* proviruses. Importantly, zip codes of high LTR-active clones were closer to enhancers and promoters than those that had low GFP+ proportions. Furthermore, these clones diminished over time in the *vpr+* pool and were disproportionately depleted in *vpr^rev^* pools, upon the addition of *vpr* to the *vpr-* pools.

We also compared the effects of LRAs on *vpr+* and *vpr-* proviral populations as well as individual clones. LRA responses vary among latency models (35), thus suggesting that combining mechanistically distinct LRAs might broaden treatment efficacy (10, 40, 59, 60). Here, when treatment with prostratin, which increases NFkB, was compared to JQ1, which enhances pTEFb availability, individual clones were affected to differing extents. Contrary to the expectation that dual treatment would enhance reactivation, prostratin plus JQ1 dual treatment suppressed some proviruses in the *vpr-* pool but did not display this phenotype in the *vpr+* pool. The suppressed proviruses tended to be relatively close to promoters and enhancers, possibly suggesting a transcriptional interference role in this effect (3, 61). The adoption by the *vpr^rev^* pools of the *vpr+* pool’s patterns of responsiveness to LRAs further corroborated the role of Vpr: even in this very limited analysis of two drugs’ effects.

Analysis here suggested a correlation between LTR-active clones and integration positions relative to certain epigenetic marks. To address the relationship of these observations to persistent populations *in vivo*, where survival during cell proliferation shapes the reservoir (62), we mapped distances to pre-existing marks for previously published integration sites of intact, inferred to be intact, or large proviruses in three patients on antiretroviral therapy (ART) (49), and compared this to integration site distances in our *vpr+* and *vpr-* pools. This analysis indicated that proximities of *vpr+* proviruses to the tested features were significantly more similar to those observed in patients than were *vpr-* pool members. This suggests that the observed effect of *vpr* in our *in vitro* system may also contribute to shaping the persistent latent reservoir in patients. If so, existing animal studies that have studied latency and reactivation using *vpr-* proviruses may largely be tracking the properties of proviruses that are eliminated during natural infection (52).

## Materials and Methods

### Tissue culture of cell lines

Two immortalized mammalian cell lines were used in this study. Human Embryonic Kidney (HEK) 293T and Jurkat cells were purchased from ATCC (Cat No. CRL3216 and TIB-152 respectively). Both cell lines were preserved as frozen stocks. 293T cells were cultured in Dulbecco’s Modified Eagle Medium (DMEM) supplemented with 10% Fetal Bovine Serum (FBS) (Gemini) and 125 μM gentamycin. Jurkat cells were cultured in RPMI media supplemented with 10% FBS, 100 U/ml penicillin, 100 μg/ml streptomycin, 2 mM glutamine and 55 μM β- mercaptoethanol. To propagate cells, frozen 1 ml aliquots of each cell line were thawed rapidly in 37° C water bath and added to 9 ml of their respective pre-warmed media. After mixing gently, cells were centrifuged at 2500 rpm for 5 minutes. Supernatant was discarded and the cell pellet was resuspended gently in pre-warmed media and plated in an appropriately sized tissue culture plate. HEK 293T cells were sub-cultured when confluency was between 75-100%. Jurkat cells were passaged such that the cell concentration was maintained between 8 x 10^5^ and 1 x 10^6^ cells/ml.

### Zip coded vector and virus production

Zip coded vector DNA templates were generated by digestion of previously described modified “inside-out” forms of GKO, which here are called GPV+ and GPV^-^ (44, 63), with ClaI and MluI. The resulting 11.4 kb DNA fragment, which was devoid of plasmid backbone, was gel purified using QIAquick Gel Extraction Kit (Cat No./ID: 28706 Qiagen, Germantown, MD). A 304 bp insert fragment was generated by PCR with with GPV^+^ or GPV^-^ as template for *vpr+* and *vpr-* respectively, Phusion High-Fidelity DNA Polymerase (New England Biolabs, Inc., Ipswich, MA), and primers 5’-GACAAGATATCCTTGATCTGNNN NNNNNNNNNNNNNNNNNGCCATCGATGTGGATCTACCACACACAAGGC-3’ and 5’- CGGTGCCTGATTAATTAAACGCGTGCTCGAGACCTGGAAAAAC-3’. The 11.4 kb and 304 bp degenerate barcode containing fragments were joined by Gibson assembly using HiFi DNA assembly mix (New England Biolabs) in a molar ratio of 1:5. 3 μg of the resulting covalently closed circular DNA was directly co-transfected with 330 ng of pHEF-VSV-G (Addgene Plasmid #22501) into 70% confluent monolayer of 293T cells in a 10 cm dish using polyethylenimine (Polysciences Inc, Warrington, PA.) at 4X total transfected DNA in 800 μl 150 mM NaCl. DMEM was replaced with 4 ml RPMI1640 medium with 10% FBS and 1% Pen/strep 24 hours post transfection and culture media was harvested by filtering through a 0.22 μm filter (Fisher Scientific. Cat. No. 09-720-511) 48 hours post-transfection.

### Generation of zip coded proviral pools

1000 μl virus-containing media was mixed with polybrene at a final concentration of 0.5 μg/ml and brought to a total volume of 2000 μl by addition of RPMI. This infection mixture was added to 5 x 10^6^ Jurkat cells and incubated in two wells of a 12-well plate at 37 C with 5% CO for 5 hours. Infected cells were then transferred to 10 ml falcon tubes and centrifuged for 5 minutes at 2500 rpm at 4 °C. Following centrifugation, supernatants were replaced with fresh media and cell pellets were resuspended and cultured in two wells of a 6-well plate at 37 °C with 5% CO_2_. At 24 hours post-infection, puromycin was added to a final concentration of 0.5 μg/ml. The infected cells were expanded into 6 cm culture plates without puromycin on day 5. Ten days post-infection, the culture supernatant was replaced with fresh media and the cultures were divided into aliquots, to be either frozen, prepared for integration-site sequencing or further expanded for four additional days and sorted into GFP+ and GFP- subpools for subsequent experiments.

### Construction of vpr+ lentiviral vector and its use

A 2933 bp in vitro DNA synthesized *IRES-vpr mKO* fragment was ordered from Genewiz (South Plainfield, NJ) using an IRES sequence from pTRIPZ-hDDX5/7 (Addgene Plasmid #71307) (64) and *vpr* and *mKO* from GPV^+^. An HIV-1 lentiviral vector fragment was generated by XbaI and MfeI digestion of pWA18puro (65) to remove its puromycin resistance cassette, and was gel purified using QIAquick Gel Extraction Kit (Cat No./ID: 28706 Qiagen, Germantown, MD). The resulting DNA fragments were joined by Gibson assembly using HiFi DNA assembly mix (New England Biolabs) following the manufacturer’s protocol to generate pEA216-1. 10 μg of pEA216-1 was then co-transfected with 5 μg of the pCMVΔR8.2 helper plasmid (Addgene Plasmid #122263) and 1 μg of pHEF-VSV-G into 70% confluent monolayers of 293T cells in a 10 cm dish using polyethylenimine (Polysciences Inc, Warrington, PA.) at 4X total transfected DNA in 800 μl 150 mM NaCl. DMEM was replaced with 4 ml RPMI1640 medium with 10% FBS and 1% Pen/strep 24 hours post transfection and culture media was harvested by filtering through a 0.22 μm filter (Fisher Scientific. Cat. No. 09-720-511) 48 hours post-transfection. Parental *vpr-* pools were infected with this filtered media and sorted 48 hours post-transduction to generate *vpr-^rev^* pools.

In determining the effect of Vpr addition on parental *vpr-* pools, zip code abundances in sequences amplified from genomic DNA of parental *vpr-* GFP+ cells and from Vpr-transduced *vpr^rev^* GFP+ cells were compared. Two categories of zip codes were defined: reduced zip codes were those that were reduced 10-fold or more after Vpr transduction, whereas those with a change of 1.5-fold or less in their relative abundances were regarded as not affected by *vpr* addition (Not reduced).

### Flow cytometry and cell sorting

For Fluorescence Activated Cell Sorting (FACS) analysis by flow cytometry, Jurkat cells were suspended in phosphate buffered saline (PBS) containing 1% FBS (FACS buffer). Dead cells were excluded from all analyses and sorting experiments using propidium iodide (PI). Acquisition was carried out on the FITC channel for GFP and PE channel for PI. Cell fluorescence was assessed using BD LRS Fortessa (BD Biosciences) and data were analyzed using FlowJo software, version 10.6 (FlowJo, LLC., Ashland, Oregon). Infected cells were sorted into GFP+ and GFP- sub- populations by flow cytometry using FACS Aria II (BD Biosciences, Franklin Lakes, NJ) or iCyt Synergy SY3200 (Sony Biotechnology, San Jose, CA) cell sorters at the flow cytometry core of the University of Michigan.

### Latency reversing agents and reactivation

JQ1 and prostratin were purchased from Sigma Aldrich. Each LRA was dissolved in DMSO (Thermofisher) to produce stocks. For each experiment, stocks were added to culture medium to achieve 2 μM JQ1 and 10 μM prostratin final concentrations. Dual LRA treatment was done by adding the two LRAs to the same culture medium to achieve their respective single LRA concentrations. For reactivation experiments, GFP- cells sorted on day 14 post-infection were co-cultured with appropriate LRA for 24 hours. Cells were then centrifuged at 2000rpm, 4°C for 5 mins, cells pellets were washed twice with ice cold FACS buffer after being stained with PI for 5 mins at room temperature. The resulting cells were then washed and assessed by flow cytometry and the p24 concentration of culture supernatants was measured by a reverse transcriptase assay.

### Zip code sequencing libraries

Zip codes were amplified from the genomic DNA of infected cells as well as from the RNA of virus released into cell media. Generation of zip coded sequencing libraries from infected cells was initiated by harvesting DNA from an aliquot of 2×10^6^ infected cells. Genomic DNA extraction was carried out using the Qiagen DNeasy Blood & Tissue Kit (Qiagen, Germantown, MD). Zip codes were then amplified by polymerase chain reaction (PCR) from 200 ng of DNA template using Phusion ® High-Fidelity (HF) DNA Polymerase (New England Biolabs) in HF Buffer. Primers were designed to flank the zip code region (primer sequences: 5‘- NNACGAAGACAAGATATCCTTGATC-3’ and 5’-NNTGTGTGGTAGATCCACATCG-3’).

Multiple copies of these primers were created, each with a unique pair of known, randomized nucleotides at the 5’ end to confirm that no cross-contamination had occurred between samples. Reactions were cycled for 29 times with 30 second extension at 72°C and a 59°C annealing temperature. Zip code amplicons were purified using the DNA Clean and Concentrator-5 kit (Zymo Research, CA. Cat. No. D4013) and eluted in 15 μL of Milli-Q ® H_2_O. To amplify zip codes from virus, a tissue culture plate of infected cells was decanted into a conical tube and centrifuged at 2500 rpm for 5 minutes. The virus-containing media was separated from the cell pellet and passed through a 0.22 μm filter. To concentrate virus, viral media was subjected to ultra- centrifugation (25,000 rpm) for 120 minutes through a 20% sucrose cushion. Viral pellets were then resuspended in 200 μl PBS and viral RNA was extracted using Quick-RNA Viral Kit (Zymo Research, CA. Cat. No. R1034 & R1035) following the manufacturer’s protocol and eluted in 10 μl RNase-free water. cDNA was synthesized using 5 μl of the eluent as template using the U3 antisense primer 5’ TGTGTGGTAGATCCACATCG-3’ and M-MLV RT RNase [H-] (Promega, WI. Cat. No. MR3681) following the manufacturer’s protocol. Zip codes were amplified from this cDNA using conditions above. The zip code amplicons were then used to generate MiSeq libraries for sequencing as described previously (44).

### Integration site sequencing

Genomic DNA was extracted from *vpr+* and *vpr-* cells 10 days post-infection using Qiagen DNeasy Blood & Tissue kit (Qiagen) and 200 ng of DNA was sheared to 1 kb fragments using Covaris M220 and micro- TUBE according to the manufacturer’s recommended settings (Covaris, Woburn, MA). HIV-1 insertion site libraries were prepared and sequenced using methods described previously (44). All sequence data has been deposited to the Sequence Read Archive (SRA) under project accession PRJNA76782.

### Quantification of virus release

Virions production was quantified using a real-time reverse-transcription PCR assay previously described by Kharytonchyk et al (65). Briefly, viral lysates were prepared by adding 5 μl of culture supernatant to 5 μl of lysis buffer. Using MS2 RNA as template, MS2 cDNA was synthesized with viral lysates and quantified by real-time PCR in one reaction. Released virus was quantified and normalized for p24 level based on values determined in parallel for reference samples.

### Zip code quantification and analysis

Zip codes were identified and quantified from Illumina sequencing reads using a previously described custom suite of tools implemented in Python (https://github.com/KiddLab/hiv-zipcode-tools). Briefly, 2x75 bp paired reads were merged together using *flash* v1.2.11 [73]. Zip codes were identified by searching for known flanking sequence (with up to 1 mismatch). Only candidate zip codes with a length of 17–23 nucleotides were considered and the read count for each unique zip code was tabulated. To identify sets of zip codes for further analysis, zip code families, which account for PCR and sequencing errors, were determined by clustering together the observed unique zip codes. Abundances for the zip codes were then determined by assigning unique zip codes to the most abundant family whose sequence was within 2 mismatches and summing their associated read counts. Only zip code families with corresponding data in the integration site data were selected for further analysis.

For each latency reversal treatment condition, clonal virus release was determined by multiplying the fractional abundances of zip codes from the cDNA sequencing libraries of each treatment by their corresponding sample’s pool p24 concentration, as measured 24 hours post LRA treatment. The resulting clonal p24 values for each zip code in prostratin, JQ1, and combination prostratin- JQ1 treated samples were divided by clonal p24 values defined for cell samples exposed only to DMSO to determine fold change after LRA treatment.

### Determination of chromatin marks

H3K27ac marks annotated for the Jurkat E-6-1 clone were sourced from http://chip-atlas.org/view?id=SRX1041803, H3K4me3 and DNase I sensitivity sites datasets were downloaded from the encode project (https://www.encodeproject.org/) with the identifiers SRX1041803, and ENCFF304GVP for genome marks pre-existing in Jurkat cells, and iENCFF341XUX, ENCFF053LHH (47). Bedtools was then used to map the distance to the closest known annotated marks. Analysis of proximity to these genome marks was done using matplotlib and scipy.stats packages in python and the results were exported into Graphpad Prism version 9.1.2 to plot graphs.

## Acknowledgements

We thank Stephen Goff, Akira Ono and Joel Swanson for helpful comments on the manuscript. This work was supported by NIH/NIAID grant number R33 AI116190 to AT and JMK and by a pilot award to AT. from the Rogel Cancer Center NIH/NCI grant number P30CA046592.

---

**S1 Fig.**
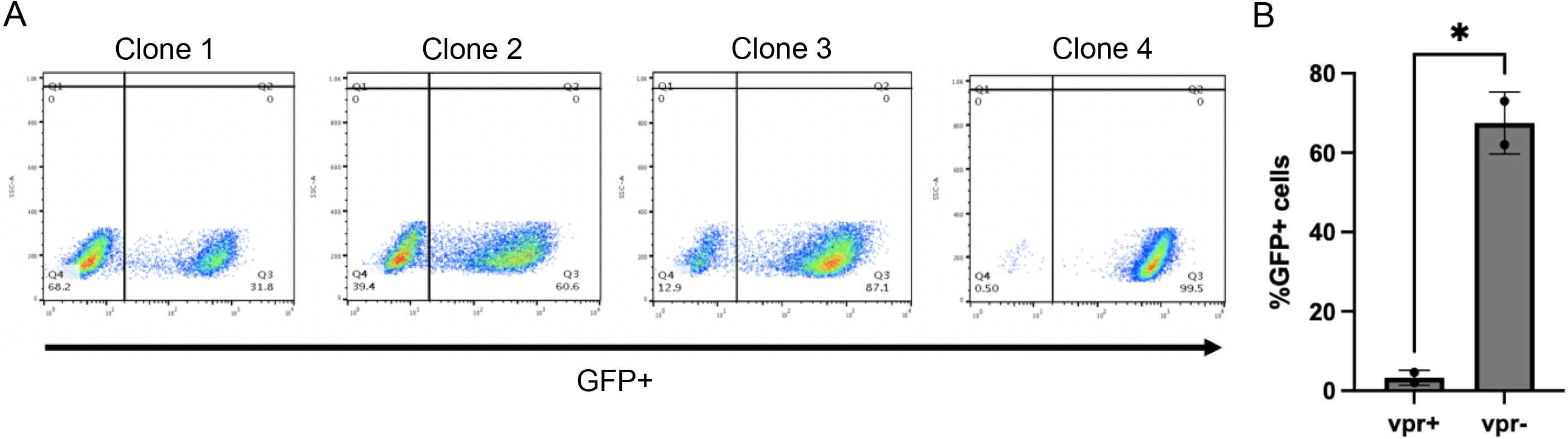
Comparison of the percent GFP+ of established cellular pools. *vpr+* and *vpr-* cell pools that were generated by infection of jurkat cells with zip coded *vpr+* and *vpr-* viruses, puromycin- selected for five days, and expanded for an additional ten days. A. Four clones were isolated by limiting dilution of *vpr-* pool, expanded for 14 days and subjected to flow cytometry. The flow plots show clonal pools 1-4 have different percent of GFP+ cells. B. Bargraph showing the percent GFP+ cells of two *vpr+* pools and two *vpr-* pools (p=* indicates p<0.05; paired t-test).

**S2 Fig.**
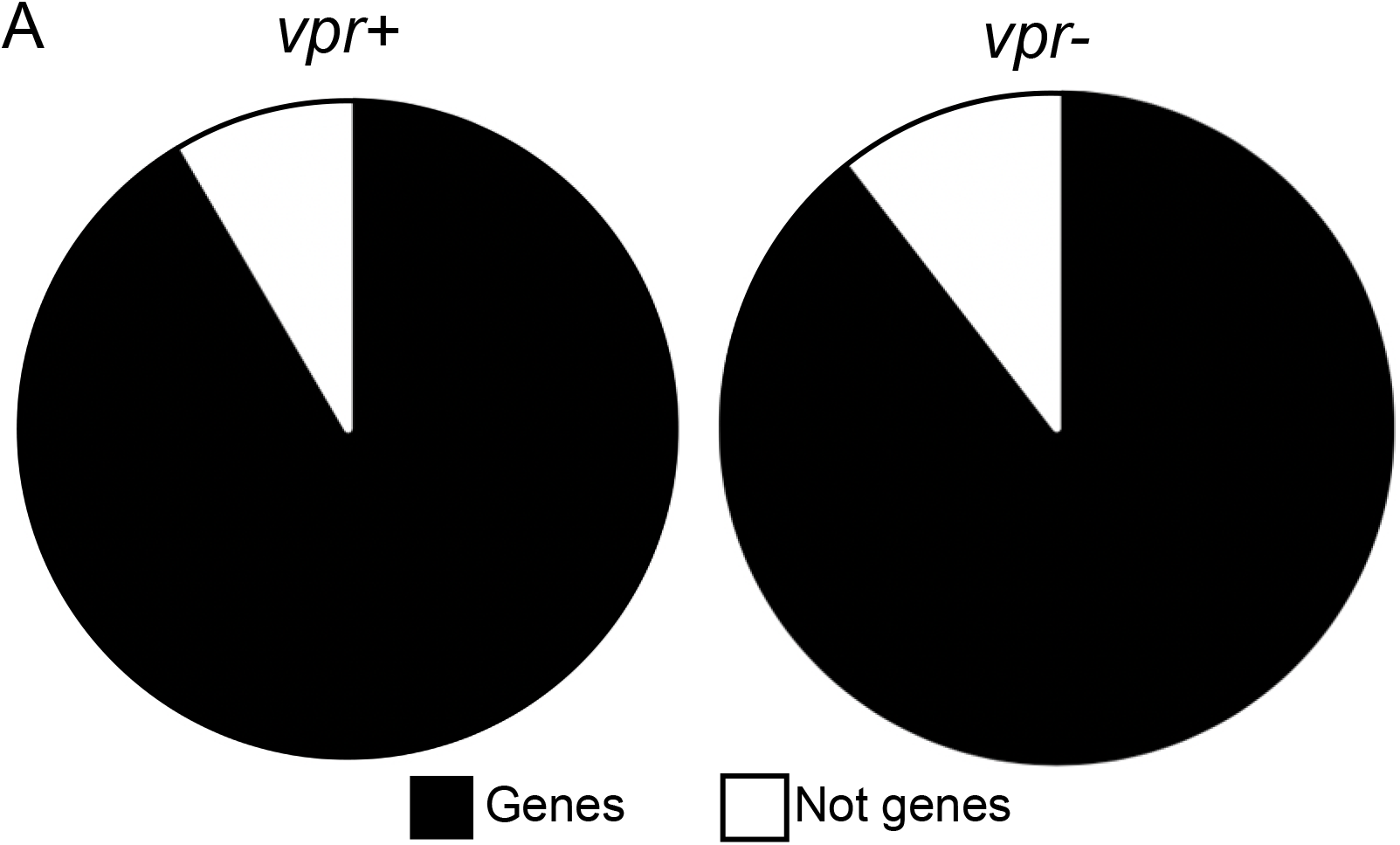
Comparison of integration sites and GFP+ proportions between *vpr-* and *vpr+* pools. A. Integration sites were sequenced and mapped to genes in Jurkat T cells. 92% of integration sites were found in genes and 8% were not in genes for *vpr*+ pool 2 (n=166) compared to 90% within genes and 10% not in genes for *vpr-* pool 2 (n=67).

**S3 Fig.**
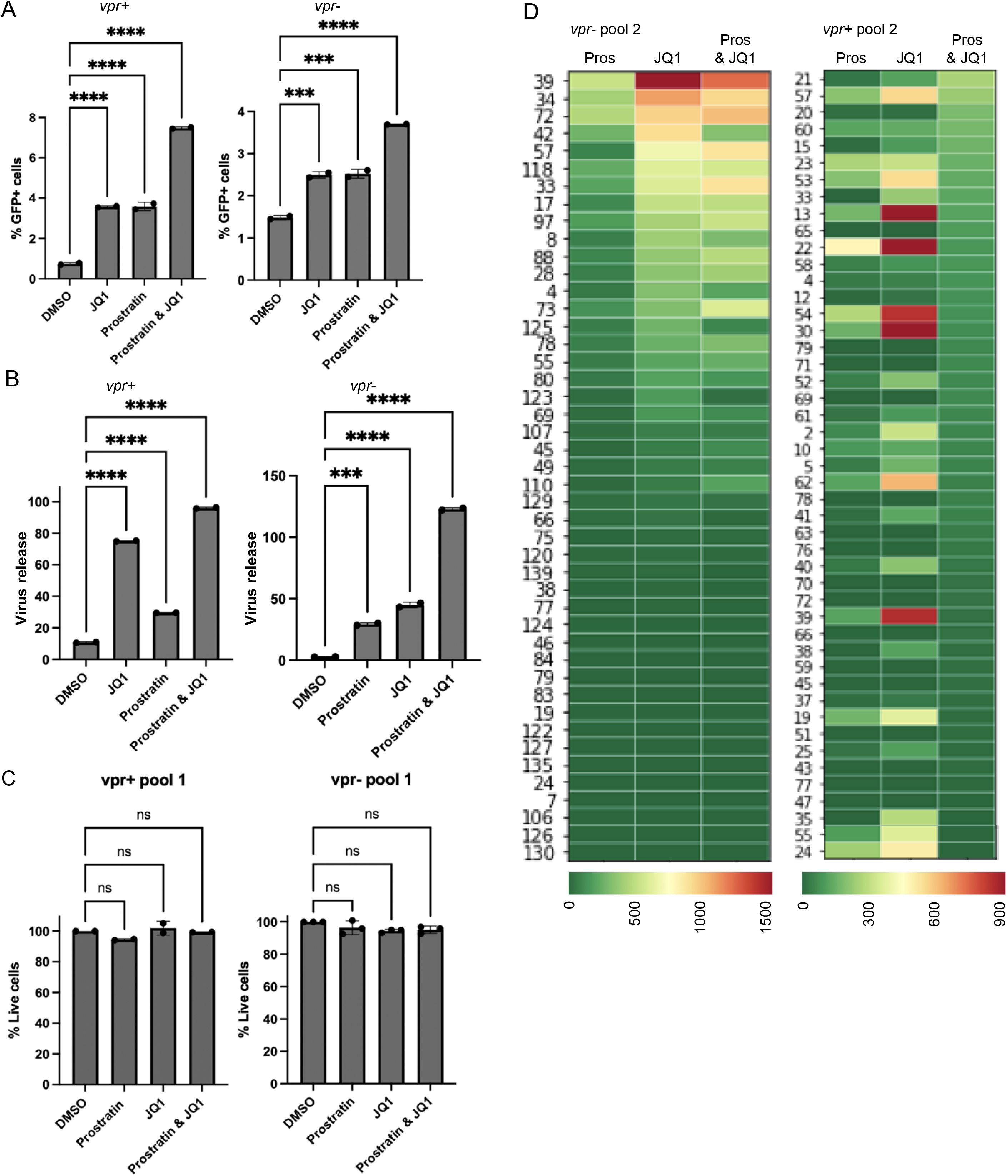
Cell viability and reactivation of additional *vpr-* and *vpr+* pools. *vpr+* and *vpr-* GFP- cells were exposed to 0.1% DMSO, 2µM JQ1, 2µM prostratin, and a combination of prostratin and JQ1. Reactivation was measured at 24 hours post-infection by flow cytometry, and virus release was quantified by reverse transcriptase assay. A. Bar graphs showing the frequency of GFP+ cells post LRA treatment (left panel for each pool) and the amount of virus released into culture supernatant (right panel for each pool) for the indicated polyclonal pools. The error bars show the mean and standard deviation from two experimental replicates. (Ordinary One-Way ANOVA; p=ns, ***, and **** indicate p>0.05, p< 0.001, and 0.0001 respectively). The error bars show mean and standard deviation. B. Viability was determined by propidium iodide staining and the percent viability was measured relative to the viability of cells in DMSO control. C. A heatmap of virus release per cell for the indicated clones (left: *vpr-* pool 2 and right: *vpr+* pool 2). Every column represents a unique cell clone’s response. The color bar indicates the extent of release in arbitrary units.

**S4 Fig.**
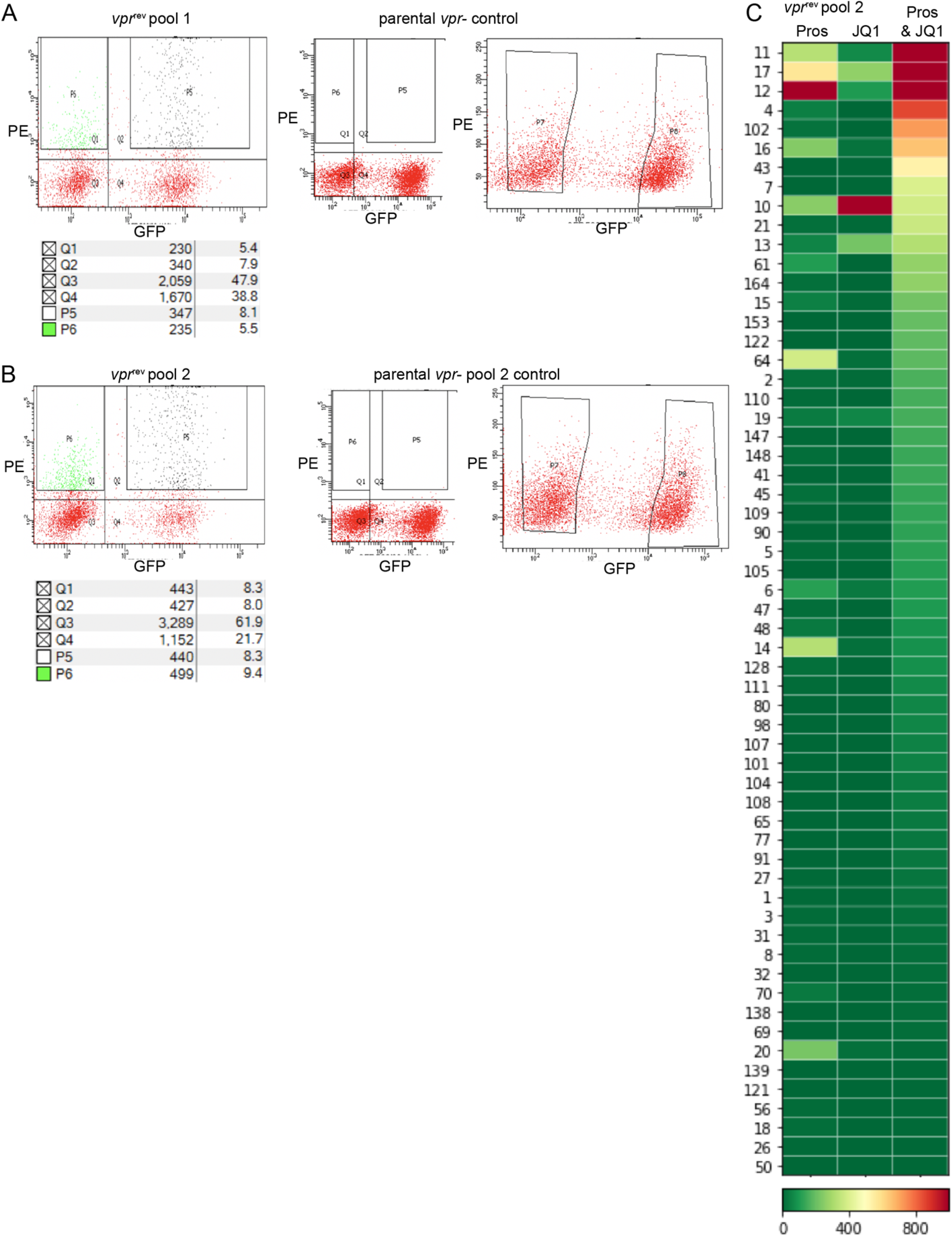
Sort gates for *vpr*-tranduced *vpr*- pools. Parental vpr- pools were transduced with Vpr- expression vector. 48 hours post-transduction, cells were sorted into PE+ GFP- and PE+ GFP+ cells. Flow plots for show the gates for sorting of A. Vpr-transduced *vpr-* pool 1 (left), parental *vpr-* pool 1 (right), B. Vpr- transduced *vpr-* pool 2 (left), and parental *vpr-* pool 2 (right). Quadrants P6 and P5 were collected for further analysis. B. Pairwise comparison of the distance to the closest H3K4me3 (left) and H3K27ac (right) marks of the zip codes that were similarly abundant at both time points (TP1 &TP2) and those that were only abundant at the first time point (TP1). Mann- Whitney two-tailed test p=ns and * indicate p>0.05 and p<0.05 respectively. C. A heatmap of virus release per treated cell for *vpr^rev^*. The indicated clones in the first column are arranged from left to right by diminishing virus release per dual treatment, and Pros means prostratin.

**S5 Fig.**
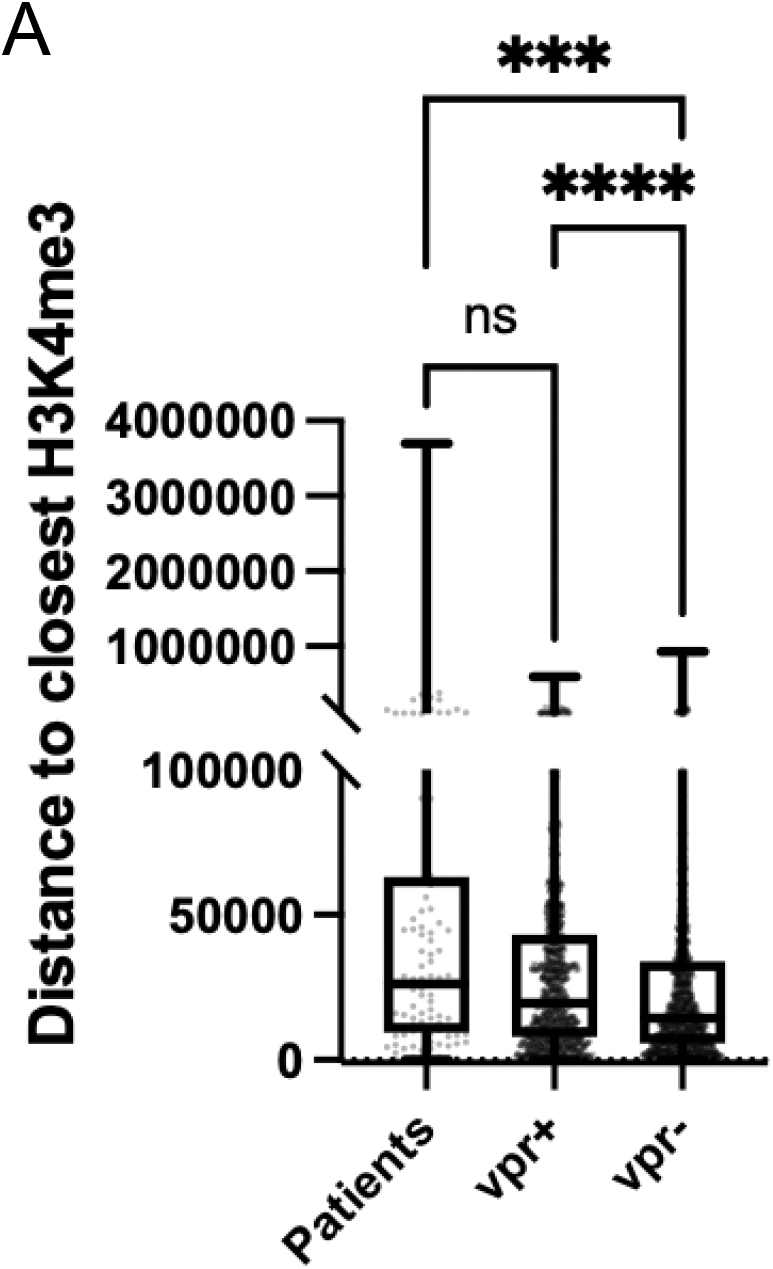
Comparison of patient integration site distance to the nearest H3K4me3 mark to *vpr+* and *vpr-* pools. Previously reported integration sites of intact proviruses from three HIV patients were mapped to the nearest pre-existing H3K4me3to in memory CD4+ T cells. Significant comparisons by Kruskal-Wallis test. p=ns, *, **, ***, **** indicates p>0.05, < 0.05, <0.01, 0.001, and 0.0001 respectively.

S Table. Table of zip codes, these clones’ fractions of abundance, and integration sites of *vpr-* and *vpr+* pools

## Notes

### Competing Interest Statement

The authors have declared no competing interest.

## References and Notes

1. Bailey J, Blankson JN, Wind-Rotolo M, Siliciano RF. Mechanisms of HIV-1 escape from immune responses and antiretroviral drugs. Current opinion in immunology. 2004;16(4):470–6.

2. Lenasi T, Contreras X, Peterlin BM. Transcriptional interference antagonizes proviral gene expression to promote HIV latency. Cell host & microbe. 2008;4(2):123–33.

3. Han Y, Lin YB, An W, Xu J, Yang H-C, O’Connell K, Dordai D, Boeke JD, Siliciano JD, Siliciano RF. Orientation-dependent regulation of integrated HIV-1 expression by host gene transcriptional readthrough. Cell host & microbe. 2008;4(2):134–46.

4. Lassen K, Han Y, Zhou Y, Siliciano J, Siliciano RF. The multifactorial nature of HIV-1 latency. Trends in molecular medicine. 2004;10(11):525–31.

5. Williams SA, Greene WC. Regulation of HIV-1 latency by T-cell activation. Cytokine. 2007;39(1):63–74.

6. Kulkosky J, Sullivan J, Xu Y, Souder E, Hamer DH, Pomerantz RJ. Expression of latent HAART-persistent HIV type 1 induced by novel cellular activating agents. AIDS research and human retroviruses. 2004;20(5):497–505.

7. Mbonye U, Wang B, Gokulrangan G, Shi W, Yang S, Karn J. Cyclin-dependent kinase 7 (CDK7)-mediated phosphorylation of the CDK9 activation loop promotes P-TEFb assembly with Tat and proviral HIV reactivation. Journal of Biological Chemistry. 2018;293(26):10009–25.

8. Archin NM, Margolis DM. Attacking latent HIV provirus: from mechanism to therapeutic strategies. Current Opinion in HIV and AIDS. 2006;1(2):134–40.

9. Verdin E, Paras P, Van Lint C. Chromatin disruption in the promoter of human immunodeficiency virus type 1 during transcriptional activation. The EMBO journal. 1993;12(8):3249–59.

10. Blazkova J, Trejbalova K, Gondois-Rey F, Halfon P, Philibert P, Guiguen A, Verdin E, Olive D, Van Lint C, Hejnar J. CpG methylation controls reactivation of HIV from latency. PLoS pathogens. 2009;5(8):e1000554.

11. Emiliani S, Fischle W, Ott M, Van Lint C, Amella CA, Verdin E. Mutations in the tat gene are responsible for human immunodeficiency virus type 1 postintegration latency in the U1 cell line. Journal of virology. 1998;72(2):1666–70.

12. Emiliani S, Van Lint C, Fischle W, Paras P, Ott M, Brady J, Verdin E. A point mutation in the HIV-1 Tat responsive element is associated with postintegration latency. Proceedings of the National Academy of Sciences. 1996;93(13):6377–81.

13. Deeks SG. HIV: Shock and kill. Nature. 2012;487(7408):439.

14. Filippakopoulos P, Qi J, Picaud S, Shen Y, Smith WB, Fedorov O, Morse EM, Keates T, Hickman TT, Felletar I. Selective inhibition of BET bromodomains. Nature. 2010;468(7327):1067–73.

15. Zhu J, Gaiha GD, John SP, Pertel T, Chin CR, Gao G, Qu H, Walker BD, Elledge SJ, Brass AL. Reactivation of latent HIV-1 by inhibition of BRD4. Cell reports. 2012;2(4):807–16.

16. Contreras X, Schweneker M, Chen C-S, McCune JM, Deeks SG, Martin J, Peterlin BM. Suberoylanilide hydroxamic acid reactivates HIV from latently infected cells. Journal of Biological Chemistry. 2009;284(11):6782–9.

17. Edelstein LC, Micheva-Viteva S, Phelan BD, Dougherty JP. Activation of latent HIV type 1 gene expression by suberoylanilide hydroxamic acid (SAHA), an HDAC inhibitor approved for use to treat cutaneous T cell lymphoma. AIDS research and human retroviruses. 2009;25(9):883–7.

18. Jiang G, Mendes EA, Kaiser P, Sankaran-Walters S, Tang Y, Weber MG, Melcher GP, Thompson III GR, Tanuri A, Pianowski LF. Reactivation of HIV latency by a newly modified Ingenol derivative via protein kinase Cδ–NF-κB signaling. AIDS (London, England). 2014;28(11):1555.

19. Archin NM, Espeseth A, Parker D, Cheema M, Hazuda D, Margolis DM. Expression of latent HIV induced by the potent HDAC inhibitor suberoylanilide hydroxamic acid. AIDS research and human retroviruses. 2009;25(2):207–12.

20. Eriksson S, Graf EH, Dahl V, Strain MC, Yukl SA, Lysenko ES, Bosch RJ, Lai J, Chioma S, Emad F. Comparative analysis of measures of viral reservoirs in HIV-1 eradication studies. PLoS pathogens. 2013;9(2):e1003174.

21. Darcis G, Kula A, Bouchat S, Fujinaga K, Corazza F, Ait-Ammar A, Delacourt N, Melard A, Kabeya K, Vanhulle C. An in-depth comparison of latency-reversing agent combinations in various in vitro and ex vivo HIV-1 latency models identified bryostatin-1+ JQ1 and ingenol-B+ JQ1 to potently reactivate viral gene expression. PLoS pathogens. 2015;11(7):e1005063.

22. Pace MJ, Agosto L, Graf EH, O’Doherty U. HIV reservoirs and latency models. Virology. 2011;411(2):344–54.

23. Jordan A, Bisgrove D, Verdin E. HIV reproducibly establishes a latent infection after acute infection of T cells in vitro. The EMBO journal. 2003;22(8):1868–77.

24. Eckstein DA, Sherman MP, Penn ML, Chin PS, De Noronha CM, Greene WC, Goldsmith MA. HIV-1 Vpr enhances viral burden by facilitating infection of tissue macrophages but not nondividing CD4+ T cells. The Journal of experimental medicine. 2001;194(10):1407–19.

25. Kino T, Gragerov A, Slobodskaya O, Tsopanomichalou M, Chrousos GP, Pavlakis GN. Human immunodeficiency virus type 1 (HIV-1) accessory protein Vpr induces transcription of the HIV-1 and glucocorticoid-responsive promoters by binding directly to p300/CBP coactivators. Journal of virology. 2002;76(19):9724–34.

26. Sato K, Misawa N, Iwami S, Satou Y, Matsuoka M, Ishizaka Y, Ito M, Aihara K, An DS, Koyanagi Y. HIV-1 Vpr accelerates viral replication during acute infection by exploitation of proliferating CD4+ T cells in vivo. PLoS pathogens. 2013;9(12):e1003812.

27. Zhang Q, Kang Y, Wang S, Gonzalez GM, Li W, Hui H, Wang Y, Rana TM. HIV reprograms host m6Am RNA methylome by viral Vpr protein-mediated degradation of PCIF1. Nature Communications. 2021;12(1):1–13.

28. Zhang F, Bieniasz PD. HIV-1 Vpr induces cell cycle arrest and enhances viral gene expression by depleting CCDC137. Elife. 2020;9:e55806.

29. Yao X-J, Mouland AJ, Subbramanian RA, Forget J, Rougeau N, Bergeron D, Cohen EA. Vpr stimulates viral expression and induces cell killing in human immunodeficiency virus type 1- infected dividing Jurkat T cells. Journal of virology. 1998;72(6):4686–93.

30. Bauby H, Ward CC, Hugh-White R, Swanson CM, Schulz R, Goujon C, Malim MH. HIV- 1 Vpr Induces Widespread Transcriptomic Changes in CD4+ T Cells Early Postinfection. mBio. 2021;12(3):e01369–21.

31. Chen H-C, Martinez JP, Zorita E, Meyerhans A, Filion GJ. Position effects influence HIV latency reversal. Nature Structural and Molecular Biology. 2017;24(1):47.

32. Jordan A, Defechereux P, Verdin E. The site of HIV-1 integration in the human genome determines basal transcriptional activity and response to Tat transactivation. The EMBO journal. 2001;20(7):1726–38.

33. Fennessey CM, Pinkevych M, Immonen TT, Reynaldi A, Venturi V, Nadella P, Reid C, Newman L, Lipkey L, Oswald K. Genetically-barcoded SIV facilitates enumeration of rebound variants and estimation of reactivation rates in nonhuman primates following interruption of suppressive antiretroviral therapy. PLoS pathogens. 2017;13(5):e1006359.

34. Jefferys SR, Burgos SD, Peterson JJ, Selitsky SR, Turner A-MW, James LI, Tsai Y-H, Coffey AR, Margolis DM, Parker J. Epigenomic characterization of latent HIV infection identifies latency regulating transcription factors. PLoS pathogens. 2021;17(2):e1009346.

35. Spina CA, Anderson J, Archin NM, Bosque A, Chan J, Famiglietti M, Greene WC, Kashuba A, Lewin SR, Margolis DM. An in-depth comparison of latent HIV-1 reactivation in multiple cell model systems and resting CD4+ T cells from aviremic patients. PLoS pathogens. 2013;9(12):e1003834.

36. Dahabieh MS, Ooms M, Simon V, Sadowski I. A doubly fluorescent HIV-1 reporter shows that the majority of integrated HIV-1 is latent shortly after infection. Journal of virology. 2013;87(8):4716–27.

37. Battivelli E, Dahabieh MS, Abdel-Mohsen M, Svensson JP, Da Silva IT, Cohn LB, Gramatica A, Deeks S, Greene WC, Pillai SK. Distinct chromatin functional states correlate with HIV latency reactivation in infected primary CD4+ T cells. Elife. 2018;7:e34655.

38. Xing S, Bullen CK, Shroff NS, Shan L, Yang H-C, Manucci JL, Bhat S, Zhang H, Margolick JB, Quinn TC. Disulfiram reactivates latent HIV-1 in a Bcl-2-transduced primary CD4+ T cell model without inducing global T cell activation. Journal of virology. 2011;85(12):6060–4.

39. Yang H-C, Xing S, Shan L, O’Connell K, Dinoso J, Shen A, Zhou Y, Shrum CK, Han Y, Liu JO. Small-molecule screening using a human primary cell model of HIV latency identifies compounds that reverse latency without cellular activation. The Journal of clinical investigation. 2009;119(11):3473–86.

40. Archin NM, Liberty A, Kashuba AD, Choudhary SK, Kuruc J, Crooks A, Parker D, Anderson E, Kearney M, Strain M. Administration of vorinostat disrupts HIV-1 latency in patients on antiretroviral therapy. Nature. 2012;487(7408):482.

41. Spivak AM, Andrade A, Eisele E, Hoh R, Bacchetti P, Bumpus NN, Emad F, Buckheit III R, McCance-Katz EF, Lai J. A pilot study assessing the safety and latency-reversing activity of disulfiram in HIV-1–infected adults on antiretroviral therapy. Clinical infectious diseases. 2013;58(6):883–90.

42. Bruner KM, Wang Z, Simonetti FR, Bender AM, Kwon KJ, Sengupta S, Fray EJ, Beg SA, Antar AA, Jenike KM. A quantitative approach for measuring the reservoir of latent HIV-1 proviruses. Nature. 2019;566(7742):120–5.

43. Wonderlich ER, Subramanian K, Cox B, Wiegand A, Lackman-Smith C, Bale MJ, Stone M, Hoh R, Kearney MF, Maldarelli F, Deeks SG, Busch MP, Ptak RG, Kulpa DA. Effector memory differentiation increases detection of replication-competent HIV-l in resting CD4+ T cells from virally suppressed individuals. PLoS pathogens. 2019;15(10):e1008074.

44. Read DF, Atindaana E, Pyaram K, Yang F, Emery S, Cheong A, Nakama KR, Burnett C, Larragoite ET, Battivelli E, Verdin E, Planelles V, Chang C-H, Telesnitsky A, Kidd JM. Stable integrant-specific differences in bimodal HIV-1 expression patterns revealed by high-throughput analysis. PLOS Pathogens. 2019;15(10):e1007903.

45. Miller CM, Akiyama H, Agosto LM, Emery A, Ettinger CR, Swanstrom RI, Henderson AJ, Gummuluru S. Virion-associated Vpr alleviates a postintegration block to HIV-1 infection of dendritic cells. Journal of virology. 2017;91(13):e00051–17.

46. ENCODE. [Available from: https://www.encodeproject.org/experiments/ENCSR000BXX/.

47. Consortium EP. An integrated encyclopedia of DNA elements in the human genome. Nature. 2012;489(7414):57–74.

48. Boehm D, Calvanese V, Dar RD, Xing S, Schroeder S, Martins L, Aull K, Li P-C, Planelles V, Bradner JE. BET bromodomain-targeting compounds reactivate HIV from latency via a Tat- independent mechanism. Cell cycle. 2013;12(3):452–62.

49. Einkauf KB, Lee GQ, Gao C, Sharaf R, Sun X, Hua S, Chen SM, Jiang C, Lian X, Chowdhury FZ. Intact HIV-1 proviruses accumulate at distinct chromosomal positions during prolonged antiretroviral therapy. The Journal of clinical investigation. 2019;129(3):988–98.

50. Kim M, Hosmane NN, Bullen CK, Capoferri A, Yang H-C, Siliciano JD, Siliciano RF. A primary CD4+ T cell model of HIV-1 latency established after activation through the T cell receptor and subsequent return to quiescence. Nature protocols. 2014;9(12):2755–70.

51. Gummuluru S, Emerman M. Cell cycle-and Vpr-mediated regulation of human immunodeficiency virus type 1 expression in primary and transformed T-cell lines. Journal of virology. 1999;73(7):5422–30.

52. Marsden MD, Zhang T-h, Du Y, Dimapasoc M, Soliman MS, Wu X, Kim JT, Shimizu A, Schrier A, Wender PA. Tracking HIV Rebound following Latency Reversal Using Barcoded HIV. Cell Reports Medicine. 2020;1(9):100162.

53. Stewart SA, Poon B, Jowett J, Chen I. Human immunodeficiency virus type 1 Vpr induces apoptosis following cell cycle arrest. Journal of Virology. 1997;71(7):5579–92.

54. Weinberger AD, Weinberger LS. Stochastic fate selection in HIV-infected patients. Cell. 2013;155(3):497–9.

55. McNamara LA, Ganesh JA, Collins KL. Latent HIV-1 infection occurs in multiple subsets of hematopoietic progenitor cells and is reversed by NF-κB activation. Journal of virology. 2012;86(17):9337–50.

56. Sherrill-Mix S, Lewinski MK, Famiglietti M, Bosque A, Malani N, Ocwieja KE, Berry CC, Looney D, Shan L, Agosto LM. HIV latency and integration site placement in five cell-based models. Retrovirology. 2013;10(1):90.

57. Poon B, Chen IS. Human immunodeficiency virus type 1 (HIV-1) Vpr enhances expression from unintegrated HIV-1 DNA. Journal of virology. 2003;77(7):3962–72.

58. Vansant G, Chen H-C, Zorita E, Trejbalová K, Miklík D, Filion G, Debyser Z. The chromatin landscape at the HIV-1 provirus integration site determines viral expression. Nucleic acids research. 2020;48(14):7801–17.

59. Rasmussen TA, Tolstrup M, Søgaard OS. Reversal of latency as part of a cure for HIV-1. Trends in microbiology. 2016;24(2):90–7.

60. Chun T-W, Engel D, Mizell SB, Ehler LA, Fauci AS. Induction of HIV-1 replication in latently infected CD4+ T cells using a combination of cytokines. Journal of Experimental Medicine. 1998;188(1):83–91.

61. Greger IH, Proudfoot NJ, Demarchi F, Giacca M. Transcriptional interference perturbs the binding of Sp1 to the HIV-1 promoter. Nucleic acids research. 1998;26(5):1294–300.

62. Reeves DB, Duke ER, Wagner TA, Palmer SE, Spivak AM, Schiffer JT. A majority of HIV persistence during antiretroviral therapy is due to infected cell proliferation. Nature communications. 2018;9(1):1–16.

63. Battivelli E, Dahabieh MS, Abdel-Mohsen M, Svensson JP, Da Silva IT, Cohn LB, Gramatica A, Deeks S, Greene WC, Pillai SK. Chromatin Functional States Correlate with HIV Latency Reversal in Infected Primary CD4+ T Cells. bioRxiv. 2018:242958.

64. Yang J, Zhao Y, Kalita M, Li X, Jamaluddin M, Tian B, Edeh CB, Wiktorowicz JE, Kudlicki A, Brasier AR. Systematic determination of human cyclin dependent kinase (CDK)-9 interactome identifies novel functions in RNA splicing mediated by the DEAD Box (DDX)-5/17 RNA helicases. Molecular & Cellular Proteomics. 2015;14(10):2701–21.

65. Kharytonchyk S, King SR, Ndongmo CB, Stilger KL, An W, Telesnitsky A. Resolution of Specific Nucleotide Mismatches by Wild-Type and AZT-Resistant Reverse Transcriptases during HIV-1 Replication. Journal of molecular biology. 2016;428(11):2275–88.

